# Spatial transcriptomic profiling of human retinoblastoma

**DOI:** 10.1101/2024.02.05.578886

**Authors:** Luozixian Wang, Sandy Hung, Daniel Urrutia-Cabrera, Roy C. K. Kong, Sandra Staffieri, Louise E. Ludlow, Xianzhong Lau, Peng-Yuan Wang, Alex W. Hewitt, Raymond C.B. Wong

## Abstract

Retinoblastoma (RB) represents one of the most prevalent intraocular cancers in children. Understanding the tumor heterogeneity in RB is important to design better targeted therapies. Here we used spatial transcriptomic to profile human retina and RB tumor to comprehensively dissect the spatial cell-cell communication networks. We found high intratumoral heterogeneity in RB, consisting of 10 transcriptionally distinct subpopulations with varying levels of proliferation capacity. Our results uncovered a complex architecture of the tumor microenvironment that predominantly consisted of cone precursors, as well as glial cells and cancer-associated fibroblasts. We delineated the cell trajectory underlying malignant progression of RB, and identified key signaling pathways driving genetic regulation across RB progression. We also explored the signaling pathways mediating cell-cell communications in RB subpopulations, and mapped the spatial networks of RB subpopulations and region neighbors. Altogether, we constructed the first spatial gene atlas for RB, which allowed us to characterize the transcriptomic landscape in spatially-resolved RB subpopulations, providing novel insights into the complex spatial communications involved in RB progression.

## Introduction

Retinoblastoma (RB) is the most common eye cancer in children under the age of 5 years, which is characterized by tumor development in the retina, resulting in blindness, loss of the affected eye, or death when disease cannot be controlled. RB affects approximately one in every ∼20,000 live births annually ^1^. While a majority of children with RB cases stem from biallelic *RB1* inactivation, for a minor portion, *MYCN* amplification is exhibited without *RB1* alterations ^2,3^. Next-generation sequencing studies have also revealed additional genetic abnormalities beyond *RB1* inactivation that are associated with aggressive histopathological features ^4,5^, highlighting the complexity of the genetic lesions causing the disease.

While genetically engineered mouse models are commonly utilized to study RB biology, they fail to fully replicate the complexities observed in human RB ^6,7^. The factors that contributed to this species-to-species variation included possible species disparities in retinal development, genetics, and tumor microenvironment at the transcriptomic, epigenetic and cellular levels ^7,8^. Recent studies using human pluripotent stem cells and retinal organoids provided a valuable *in vitro* model to study the cellular abnormalities caused by biallelic mutations of the *RB1* ^9,10^. In addition, to study the cellular heterogeneity of RB, a number of recent studies have profiled primary RB tumors using single cell RNAseq (scRNA-seq)^5,11,12^ , advancing our understanding of the cellular origin of RB and the pathological mechanisms driving RB. However, these single cell transcriptomic datasets do not retain structural information which would be important to study the collective tissue architecture of RB.

Spatial transcriptomics provide an exciting opportunity to address this, by simultaneous profiling of gene expression and their spatial distribution in the tissue. Notably, spatial transcriptomic profiling has yielded critical insights into tumor heterogeneity, interactions within the tumor microenvironment (TME) and identified potential therapeutic targets in melanoma and oral squamous cell carcinoma ^13,14^. Here we present the first report of a spatial gene atlas for human RB using spatial transcriptomic profiling, providing detailed spatial perspective of transcriptome alteration underlying retinal cancer. Our results identified distinct RB subpopulations and highlighted the spatial communication networks in RB. These findings not only deepen our understanding of RB biology but also highlight the versatility of spatial transcriptomics in unraveling complex molecular architecture in tumors.

## Results

### Spatial transcriptomic profiling of human RB and control retina

Using 10X Visium, we first performed tissue optimization assay on human control retina to determine the optimal permeabilization time for retinal tissues (Supplementary figure 1). Next, we performed spatial transcriptomics on 3 control retinas (n=4 tissue sections) and a RB tumor sample from a 4 year old patient (n=4 tissue sections; Figure 1A and Supplementary table 1). After pre-processing and quality control, in the control retina samples we manually annotated the cell types in the captured regions, corresponding to retina, choroid, sclera and optic nerve (total 3761 spots, Figure 1B). In the RB sample, we captured 2982 spots and detected 16398 unique genes (Figure 1B). Using non-linear dimensionality reduction, we identified ten transcriptionally distinct clusters in RB (C0-C9, Figure 1C).

**Figure 1.**
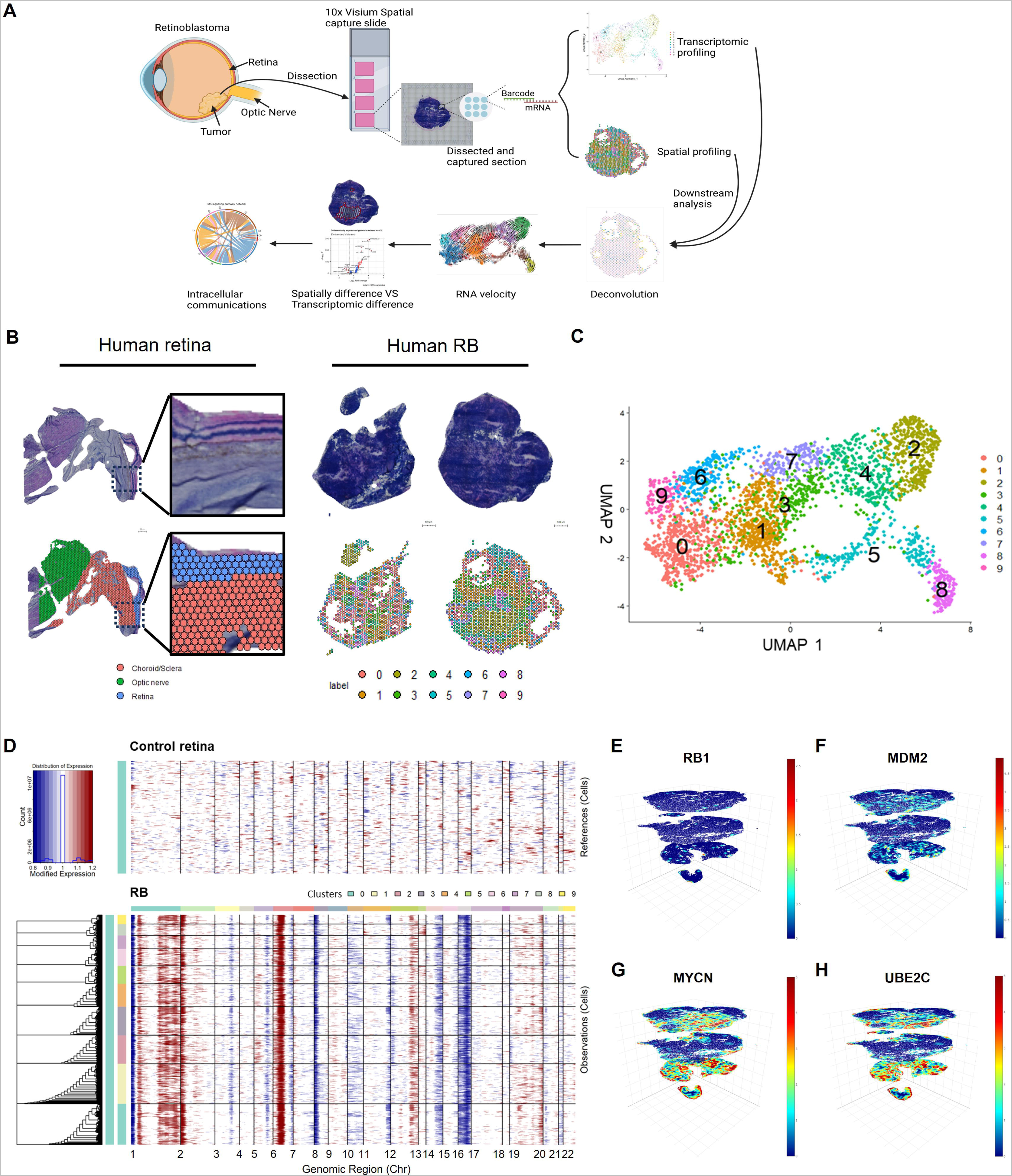
Spatial transcriptomic profiles of human retina and RB. A) Overview of the spatial analysis workflow to profile RB sections and downstream bioinformatics analysis pipeline. B) Visualization of the morphology of human retina and RB sections by H+E staining (top) and the spatial distribution of UMAP clusters (bottom). C) UMAP dimensional reduction with Harmony embeddings identified 10 transcriptionally distinct clusters in RB. D) Chromosomal landscape of inferred CNVs based on RB spatial transcriptomics and normal human retina spatial transcriptomics. The x-axis shows the chromosome region and the y-axis represents the samples. The colors in the heatmap represent the CNV gain (red) or loss (blue). 3D Feature plot showing expression of RB-associated genes E) *RB1*, F) *MDM2*, G) *MYCN* and H) *UBE2C*.

To study the genotype of the RB sample, we inferred copy number variations (CNVs) for each cell by averaging relative expression levels across intervals of the genome and calculated the CNV mean square in the control retina and RB samples ^15,16^. We found various CNV types in the RB sample, including gain of chromosome (Chr) 1q, 6p and loss of Chr 16 (Figure 1D), consistent with previous studies in human RB ^5^. Also, CNV patterns were also conserved across all ten RB clusters. Analysis of known RB-associated oncogenes and tumor suppressor genes revealed that *RB1* was absent in almost all parts of the RB tumor (Figure 1E, supplementary figure 2). On the other hand, *MDM2* and *MYCN* were highly expressed in many regions of the RB tumor, which play important roles in RB proliferation and tumor formation ^5,17^ (Figure 1E). Also, the majority of the RB tumor expressed *UBE2C*, a prognostic marker associated with malignant progression of RB tumor ^11^ (Figure 1E).

### Cone precursors represent the dominant population in human RB

For cell annotation in the RB clusters, we evaluated the expression level of a panel of canonical markers for major retinal cell types and calculated a ‘Cell score’ for individual RB clusters (Supplementary table 1). All ten RB clusters exhibited low scores for retinal progenitor cells, pigment epithelium cells, rod cells, horizontal cells, amacrine cells, and retinal ganglion cells (Figure 2A). C1-8 demonstrated high expression of cone cell markers, with a similar but less pronounced trend observed for markers of bipolar cells.

**Figure 2.**
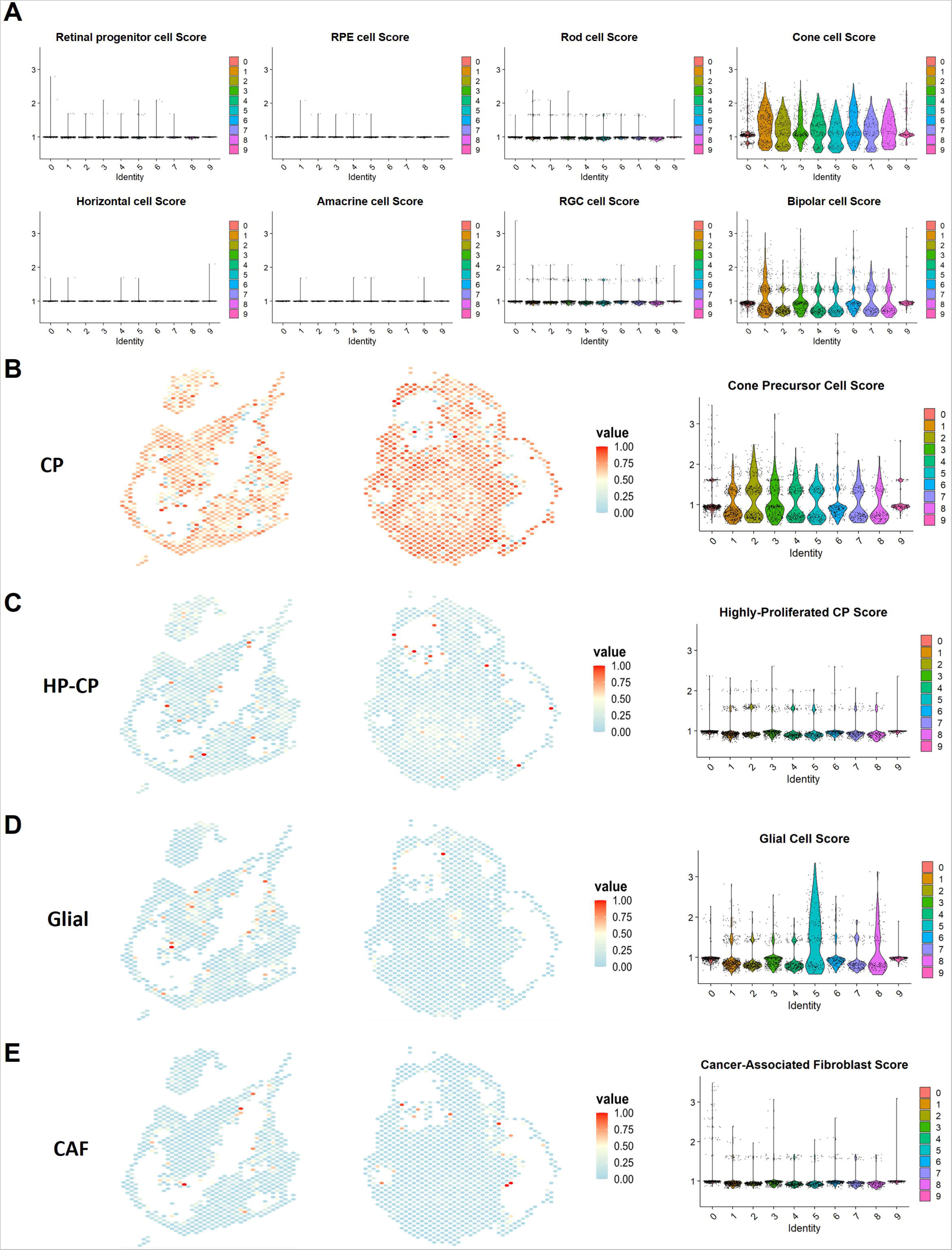
Cell type annotation using marker score and deconvolution. A) Violin plots of the aggregated expression of reference marker gene sets across RB clusters (C0-C9). The X-axis represents each RB cluster and the Y-axis indicates the averaged expression level of reference retinal markers. RPE: retinal pigmented epithelium; RGC: retinal ganglion cells. Cell deconvolution identified spatial distribution of B) cone precursors (CP), C) highly-proliferated cone precursors (HP-CP), D) glial cells and E) cancer-associated fibroblast (CAF) within RB. Right panel: Violin plots showing cell scores of respective marker genes in the identified RB clusters (C0-C9).

Considering that the resolution of spatial transcriptomic profiling using the Visium platform can not reach true single cell level in retinal cells, we perform cell-type deconvolution to resolve cell type composition in our RB sample using a reference RB single-cell RNAseq dataset ^5^. For deconvolution, we utilized the CARD algorithm which achieved one of the best performance in benchmarking studies ^18,19^. Notably, our deconvolution result revealed that the majority of the RB sample consisted of cone precursors (CPs), which were widely distributed in C1-C4 as well as C7-C8 (Figure 2B, Table 1). The highly-proliferative cone precursors (HP-CPs) previously defined as a more malignant state of CPs with high expression of proliferation-related genes including MKI67, TOP2A and KIF14, were present as a minor cell type in various RB clusters, such as C2, C4 and C5 (Figure 2C, Table 1). We also detected glial cells in C5 and C8 cells, as well as cancer-associated fibroblasts (CAFs) in C0, C6 and C9 (Figure 2D-E, Table 1). We further verified the deconvolution result by extracting the gene signatures of CP, HP-CP, glial and CAF from the reference scRNA-seq dataset, and confirming the expression level of these marker genes in our spatial dataset (Figure 2B-E). Previous scRNA-seq studies showed that some RB cells can retain hybrid gene expression signatures of multiple cell types ^11^, which could explain the reason that certain clusters (e.g. C8) display scores for multiple cell types. Altogether, our results support the CP origin of RB, with pathological changes in the CPs and HP-CPs being involved in human RB tumorigenesis consistent with previous report ^5,20^.

**Table 1:** Cell annotation using probability scores for cell types within clusters, as calculated by the CARD deconvolution algorithm. CP: cone precursors; HP-CP: Highly proliferative cone precursors; CAF: cancer-associated fibroblasts.

| Cluster | CP | HP-CP | Glial cell | CAF | Rod | Cone | Dominant cell type | Minor/associated cell type |
| --- | --- | --- | --- | --- | --- | --- | --- | --- |
| C0 | 0.55 | 0.14 | 0.06 | 0.12 | 0.04 | 0.09 | CP | HP-CP/CAF |
| C1 | 0.69 | 0.11 | 0.03 | 0.06 | 0.04 | 0.07 | CP | HP-CP |
| C2 | 0.62 | 0.23 | 0.03 | 0.07 | 0.01 | 0.04 | CP | HP-CP |
| C3 | 0.67 | 0.14 | 0.03 | 0.07 | 0.03 | 0.06 | CP | HP-CP |
| C4 | 0.69 | 0.16 | 0.03 | 0.05 | 0.02 | 0.05 | CP | HP-CP |
| C5 | 0.64 | 0.16 | 0.10 | 0.04 | 0.02 | 0.04 | CP | HP-CP/Glial cell |
| C6 | 0.60 | 0.13 | 0.05 | 0.12 | 0.04 | 0.06 | CP | HP-CP/CAF |
| C7 | 0.72 | 0.14 | 0.03 | 0.04 | 0.02 | 0.05 | CP | HP-CP |
| C8 | 0.61 | 0.10 | 0.18 | 0.03 | 0.03 | 0.05 | CP | HP-CP/Glial cell |
| C9 | 0.49 | 0.14 | 0.05 | 0.20 | 0.04 | 0.08 | CP | HP-CP/CAF |

### Heterogeneous RB sub-populations with varying proliferative potential

Subsequently, we analyzed the cell cycle stage and proliferative ability of cells in the samples which is an important aspect of tumor development and progression ^21^. We analyzed the expression pattern of S phase markers *PCNA* and *MCM6*, and G2/M phase markers *TOP2A* and *MKI67*, across all RB clusters (Figure 3A). Next we calculated a cell cycle score based on expression of these cell cycle markers. Interestingly, we observed that the C2 cells showed the highest G2/M scores, while other clusters like C1 and C6s showed high S scores (Figure 3B). As expected, about 97.3% cells in the RB tumor are proliferative, compared to control retina cells which are non-proliferative (Supplementary figure 3). Our results highlighted differential proliferation potential in the RB clusters, with ∼C2 cells having the highest proliferation potential (about 50.2% of cells in G2/M phase), while C8 cells have the least proliferation potential (about 85.8% cells in G1 phase). (Figure 4C). Altogether, the ten RB clusters have different proliferative capacity, representing cell subpopulations with varying malignancy within the RB sample.

**Figure 3.**
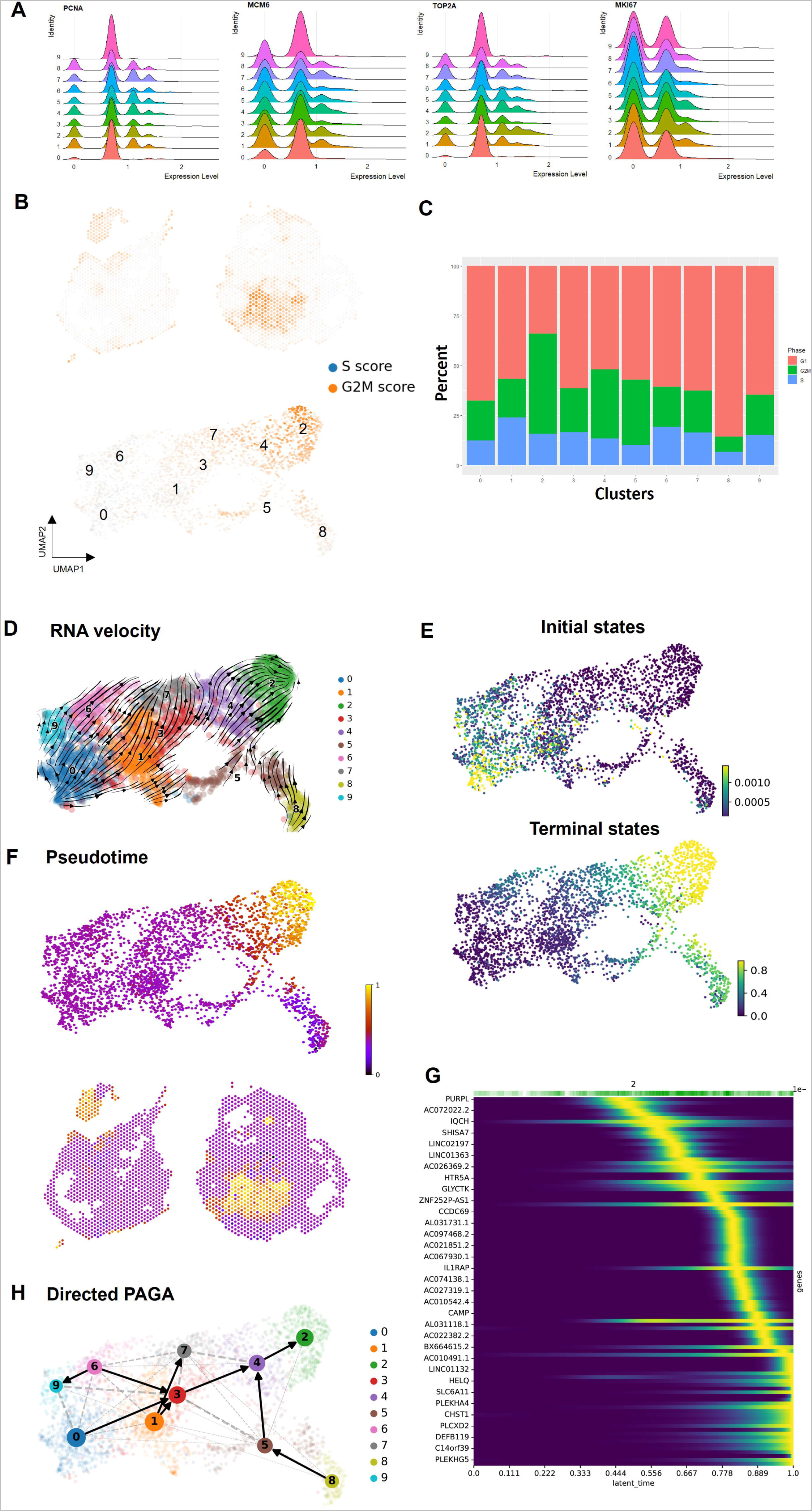
Cell-cycle and trajectory analysis of human RB. A) Ridgeline plot of the distribution of cell cycle markers across RB clusters, including S phase markers (*PCNA*, *MCM6)* and G2/M phase markers (*TOP2A*, *MKI67*). B) Cell cycle scores projected onto spatial coordinates of tissue section (top panel) and UMAP plot for RB clusters (bottom panel). C) Bar chart showing the ratio of cells in different cell cycle phases in RB clusters. D) RNA velocity projected onto UMAP. The arrows or the directional flows specify the velocity vectors and relative positions. E) Initial state prediction projected onto UMAP. The initial spots of the progression are automatically selected from the set of macrostate via a stability criterion. Terminal state prediction projected onto UMAP. The terminal spots of the progression are automatically selected from the set of macrostate via a stability criterion. F) Pseudotime projected onto UMAP (top panel) and spatial coordinates (bottom panel). Scale indicates the average number of steps cells take to reach the final state along the directional flows from the initial state. G) Top genes of the lineage drivers along the progression towards the terminal state. Driver genes involved in fate choice were plotted in a heatmap along the lineage. H) Partition-based graph abstraction (PAGA) projected onto UMAP benchmarking trajectory inference. PAGA is directionalized by velocity vectors and fate choice, providing the progression topology with connectivity between two clusters.

**Figure 4.**
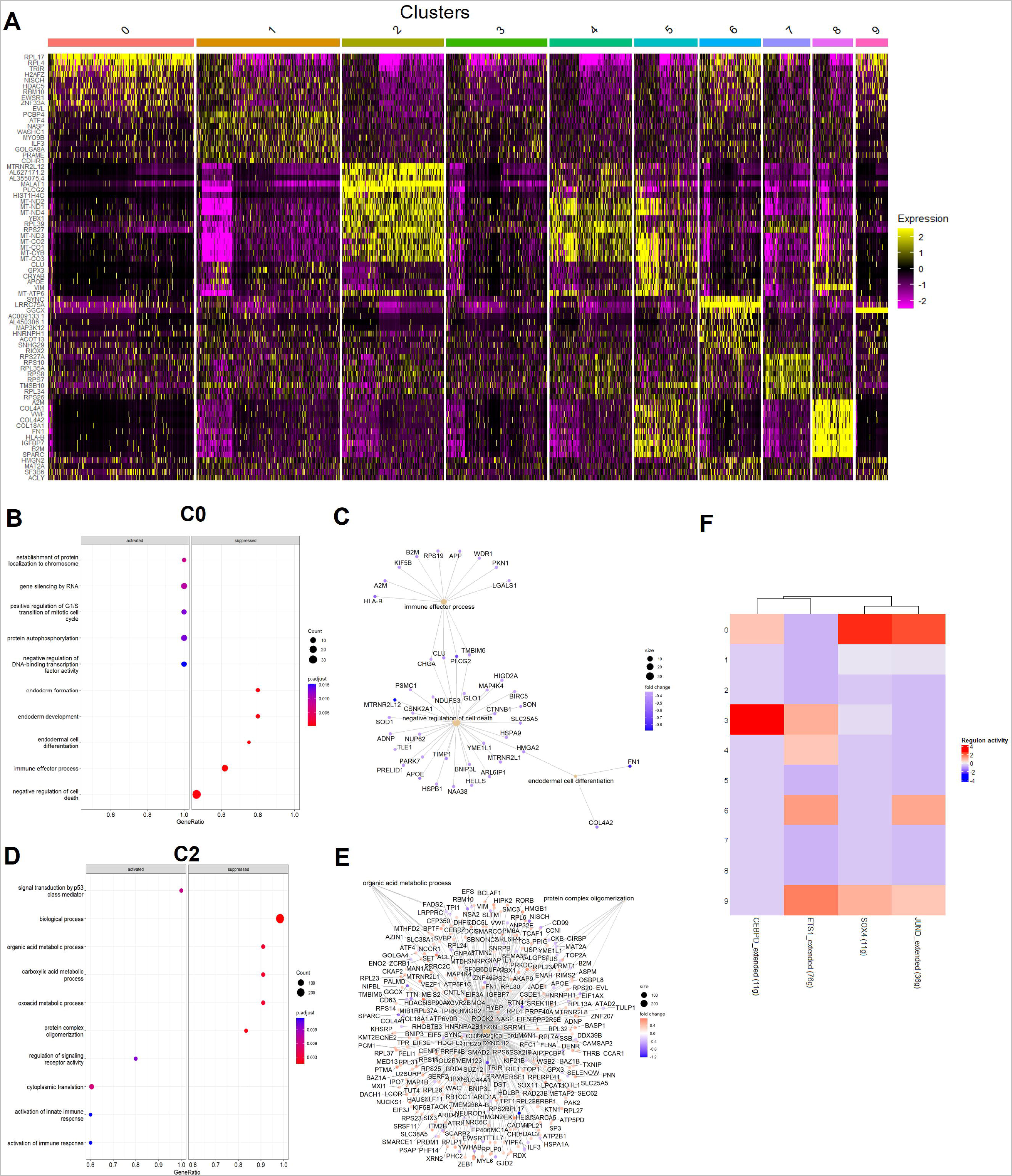
Molecular signatures and reglons in RB subpopulations. A) Heatmap showing the top10 markers which define the cluster across all ten UMAP clusters. B-E) Dot plot and category net plot showing the top GO Biological Process terms enriched in B-C) the initial state C0 and D-E) the terminal state C2. F) Heatmap showing the average regulon activity by cluster. The “extended” regulons include motifs that have been linked to the TF by motif similarity as an extension of high confidence annotations such as direction annotation or orthological inference.

### Understanding tumor progression in RB

To understand the tumor progression stages, we performed cell trajectory analysis for the RB clusters. RNA velocity was used to predict the lineage and transition between clusters by estimating the steady-state-ratio of unspliced (pre-mature) to spliced (mature) mRNAs ^22^. Our results identified root nodes in C0, C1, C8, and C9, as well as terminal nodes in C2 and C7, representing various stages in RB tumor progression(Figure 3D). Next, CellRank analysis was used to learn directed probabilistic state-change trajectories from the directional RNA velocity information to calculate fate probabilities and uncover putative lineage drivers plus lineage-specific gene expression trends ^23^. We showed that C0 and C9 cells were identified as the initial state and C2 cells were the terminal state during RB progression, while other clusters including C3 and C4 represented intermediate states (Figure 3E). Consistent with this, our inferCNV analysis also showed that C0 cells had relatively low CNV scores among the RB clusters, while C2 cells showed high CNV scores, which correlate with their roles in varying stages of RB trajectory (Supplementary figure 4).

In addition, we performed pseudo-time analysis to measure the trajectory distance of each cell transitioning between stages of RB progression ^24^. Following assignment of cells according to their position in the pseudotime trajectory, putative gene expression cascades specific to the C2 fate commitment were computed (Figure 3F). *PURPL* was identified as a key regulator among the top genes driving the initial state of RB, consistent with its function in activating p53-associated mitosis ^25^ (Figure 3G). In contrast, immune-associated genes like *HTR5A* and *ZNF252P-AS1* drive later stages of RB, both of which were shown to play a key role in regulating tumor growth ^26,27^. Also, cancer biomarkers such as *PLEKHA4* that facilitates Wnt/ß-catenin-mediated proliferation in glioblastoma, and *PLEKHG5* that promotes tumor cell migration, invasion, and angiogenesis, were also identified as key drivers of RB progression^28,29^.

To understand the global relationship between RB clusters, we computed a partition-based graph abstraction (PAGA) graph, which showed a RB trajectory connecting C0 as the initial state to intermediate states C3 and C4, followed by progressing to the terminal state C2 ^30^ (Figure 3H). We also identified two additional minor lineages that involved CAFs (C6 to C9) and glial cells (C8 to C5), supporting their roles to form the TME in RB progression. Taken together, our results highlighted a tree-like cell trajectory underlying RB progression from early to terminal stages, providing insights into the interaction of different retinal cell types that contribute to the heterogeneity in the TME of RB.

### Molecular signature of RB subpopulations

Markers that define individual RB clusters were identified to further understand the molecular mechanism in the RB subpopulations (Figure 4A). Overall, the gene signature of each cluster is consistent with their role during cellular lineage. C0 cells, representing the initial state, expressed marker genes involved in cancer proliferation like *RPL17*, and its regulator *H2AFZ* ^31,32^. Gene Ontology (GO) analysis highlighted that the C0 marker genes are involved in cell cycle regulation, along with the suppression of immune activities and cell death (Figure 4B-C). During the intermediate stages of RB we observed expression of genes involved in cell adhesion like *EVL* and *CDHR1* ^33,34^ as well as genes in MT-CO and MT-ND families ^35,36^ (C4, C5) which indicate increased oxidative phosphorylation and ATP production in RB progression (Figure 4A). GO term analysis indicated elevated protein synthesis in the intermediate state (C4, Supplementary figure 5), which is important in tumor progression ^37^. Similarly, C7 cells also showed high ribosomal protein gene expression, indicating increased translation in the tumor cells. The high expression of *VIM* confirmed the glial cell composition of the C8 cluster, and their signature of collagen genes, *VWF* and *FN1* emphasized their role in cell adhesion, migration, and differentiation ^38–40^.

On the other hand, C2 cells, representing the terminal state of RB progression, shared a similar gene signature with the preceding C4 cluster, such as the high expression of the MT-CO and MT-ND families, supporting a high growth phenotype consistent with their malignant tumor cell state (Figure 4A). Additionally, several cancer-associated genes also emerged in the C2 molecular signature, including *MRTNR2L12, MALAT1* and *YBX1.* Furthermore, GO analysis in C2 revealed gene enrichment involved p53 signaling as well as activation of immune responses (Figure 4D-E). Altogether, the elevated expression of prognosis markers or oncogenes in C2, further support that it represents a more advanced stage of tumor progression in our RB sample.

### Regulons driving RB progression

Next, we explored the gene regulatory networks in the RB subpopulations. We utilized single-cell regulatory network inference and clustering (SCENIC) to reconstruct gene regulatory networks and identify upstream regulators that establish transcriptional states and progress through state transitions ^41^. SCENIC analysis identified four regulons in the RB clusters, namely *JUND*, *SOX4*, *ETS1*, and *CEBPD* (Figure 4F, Supplementary table 2). *JUND* is known for its role in promoting cancer cell proliferation and angiogenesis ^42^ and its regulon consists of 12 high-confident transcription factors (TFs) and 36 targeted genes. *ETS1* is associated with immune response regulation and aggressive tumor progression ^43^ and its regulon comprised of 25 direct-binding TFs and 76 targeted genes. Also, *CEBPD* ^44^ and *SOX4* ^45^ are involved in tumor cell differentiation and contain 11 targeted genes in their regulons.

The regulon score heatmap across all RB clusters highlighted their involvement in transition of transcriptional states and supported the predicted RB cell trajectory (Figure 4F). Interestingly, C0 showed the highest *JUND* and *SOX4* regulon scores, indicating that tumorigenesis at the early stages is facilitated by the putative TFs in these regulons. As RB progresses, *ETS1* and *CEBPD* regulons which are involved in immune response regulation become activated in C3 and C4 intermediate state cells. In contrast, the terminal state in RB is marked by downregulation of all 4 regions, as evidenced by the low regulon scores in C2.

### Spatial networks of RB subpopulations and region neighbors

Spatial transcriptomics can dissect the spatial network within the cells in the tissue section, allowing us to perform neighborhood analysis to characterize the intercellular interactions with neighboring cells in the RB clusters. Using C0 and C2 cells as the initial and terminal states respectively, we defined the connected spatial networks by selecting all region neighbors adjacent to the specific cluster, and extracted the gene signatures of these neighboring cells to determine intercellular interaction based on spatial construction (Figure 5A-B). From differential expression (DE) analysis of C0 and its neighbors, we showed that the ribosomal proteins *RPL17* and *RPL4* were upregulated in C0, while *GGCX* and *LRRC75A* were significantly upregulated in C0 neighbors (Figure 5C). *GGCX* is a metabolic gene associated with retinitis pigmentosa and pseudoxanthoma elasticum-like skin manifestations ^46^. *LRRC75A* is involved in VEGF secretion of mesenchymal stromal cells ^47^. Interestingly, *LRRC75A* and *GGCX* were identified as the markers for C6 and C9, both of which contained CAFs, potentially supporting a role of neighboring CAFs in the TME in early stages of RB progression.

**Figure 5.**
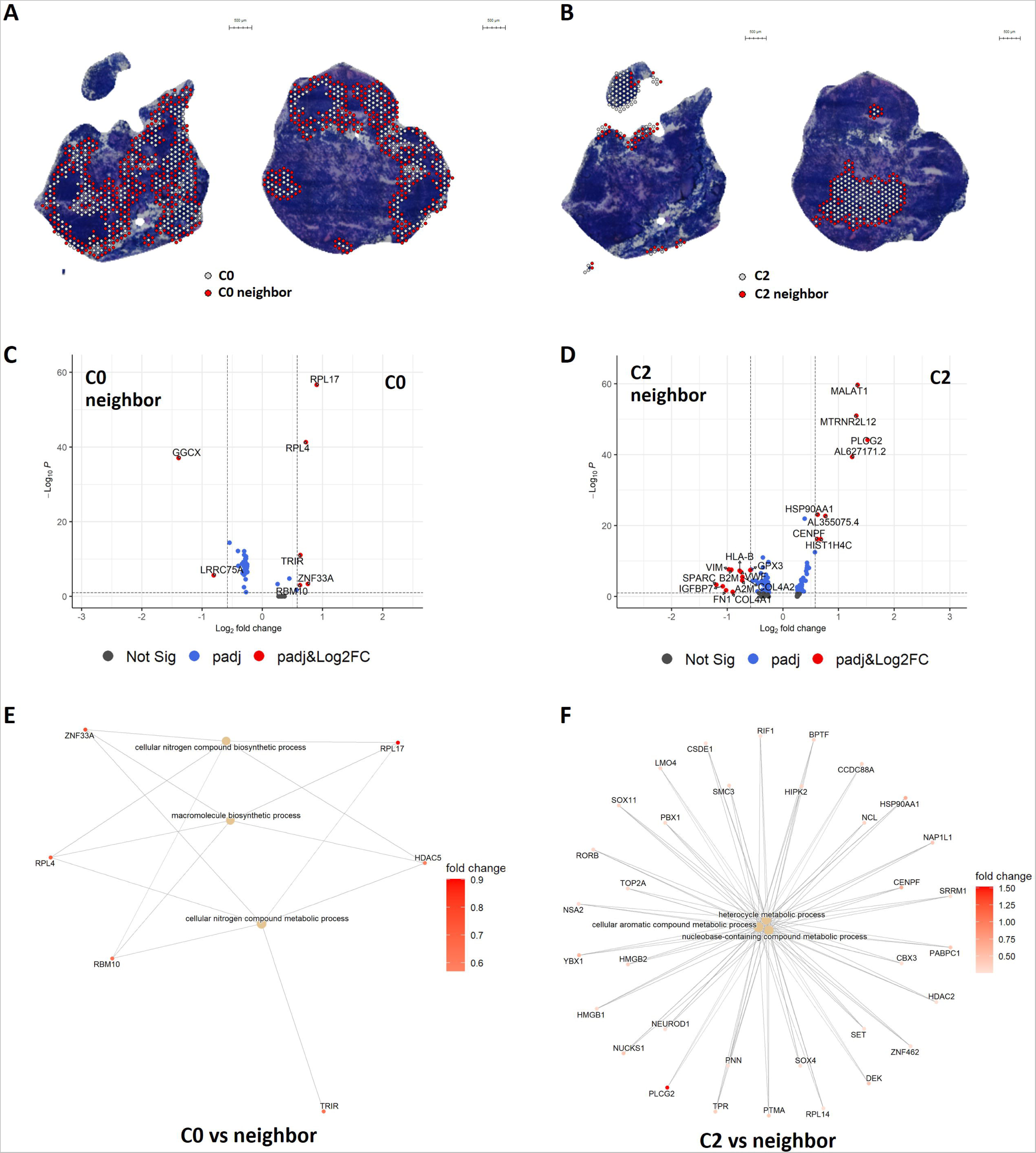
Spatial networks of RB subpopulations and region neighbors. A-B) Spatial identification of RB cells in the initial state (C0) and the terminal state (C2) with their respective region neighbors. C-D) Volcano plot showing the differential expression pattern with region neighbors for C0 and C2. E-F) Category net plots depict the linkages of genes and GO Biological Process network for C0 and C2 with their respective neighbors.

On the other hand, cancer associated genes *PLCG2*, *MALAT1* and *MTRNR2L12* were identified as the top upregulated DE genes in C2 (Figure 5D). Top DE genes for C2 neighbors included genes involved in fibronectin signaling (*FN1*), IGF signaling (*IGFBP7*) and mediators of tumor-stromal interactions (*SPARC, COL4A1*). Also we performed enrichment analysis for DE genes in C0 neighbors and C2 neighbors (Figure 5E-F). Interestingly, we found that the biosynthetic and metabolic processes to be prominent in both C0 and C2 neighbors, supporting a key role of metabolic changes in neighboring cells that contribute to RB progression.

### Cellular communication networks in human RB subpopulations

Next, we utilize CellChat ^48^ to investigate genes for signaling ligands, receptors, their cofactors and systematically analyze the cell-cell communication in RB. Our results highlighted a complex aggregated cell-cell communication network between RB subpopulations (Figure 6A). In total our results identified 25 significant signaling pathways across the ten RB clusters, many of them were associated with C8 cells. These pathways were classified into four groups based on the functional similarity of the cellular networks (Figure 6B). Specifically, Group 1 encompassed *ANGPT*, *PDGF*, *CDH5*, *FN1*, *PECAM1*, *ESAM*, *GRN*, *SPP1* and Collagen signaling pathways, and are implicated in promoting tumor growth through angiogenesis, immune evasion, abnormal cell proliferation and differentiation ^49–57^. Group 2 consisted of *MIF*, *VEGF*, *CD46* and Laminin signaling pathways, which are associated with cell adhesion and migration to support metastasis ^58–61^. Group 3 included *NOTCH*, *CD99*, *NCAM*, *JAM*, *CADM*, *MK* and *CDH* signaling pathways, with roles in tumor progression and invasion by regulating cell proliferation, apoptosis, adhesion and migration ^62–68^. The *MZP* signaling pathway is an exception in this group as it is involved in myelin sheath formation and maintenance in the nervous system. The role of *MZP* signaling in cancer remains unclear, though limited studies suggested its involvement in inducing cell death ^69^, consistent with the common characteristics of the other Group 3 pathways. Group 4 represented *NECTIN*, *SEMA5*, *APP* and *SEMA6* signaling pathways which are involved in cell migration and axon guidance ^70–73^. Our analysis highlighted complex signaling networks between the RB clusters. For instance, C8 is the primary receiver for signals from other RB clusters via the *FN1*, *VEGF* and *NOTCH* pathways (Figure 6C). In the latter case for *NOTCH* pathway, a number of clusters were also identified as mediators (eg C7) and influencers (eg. C5, C7, C9) of this signaling communication. On the other hand, C5 received signals via the *APP* pathways from a range of senders (eg. C8) and influencers (eg. C7, C8).

**Figure 6.**
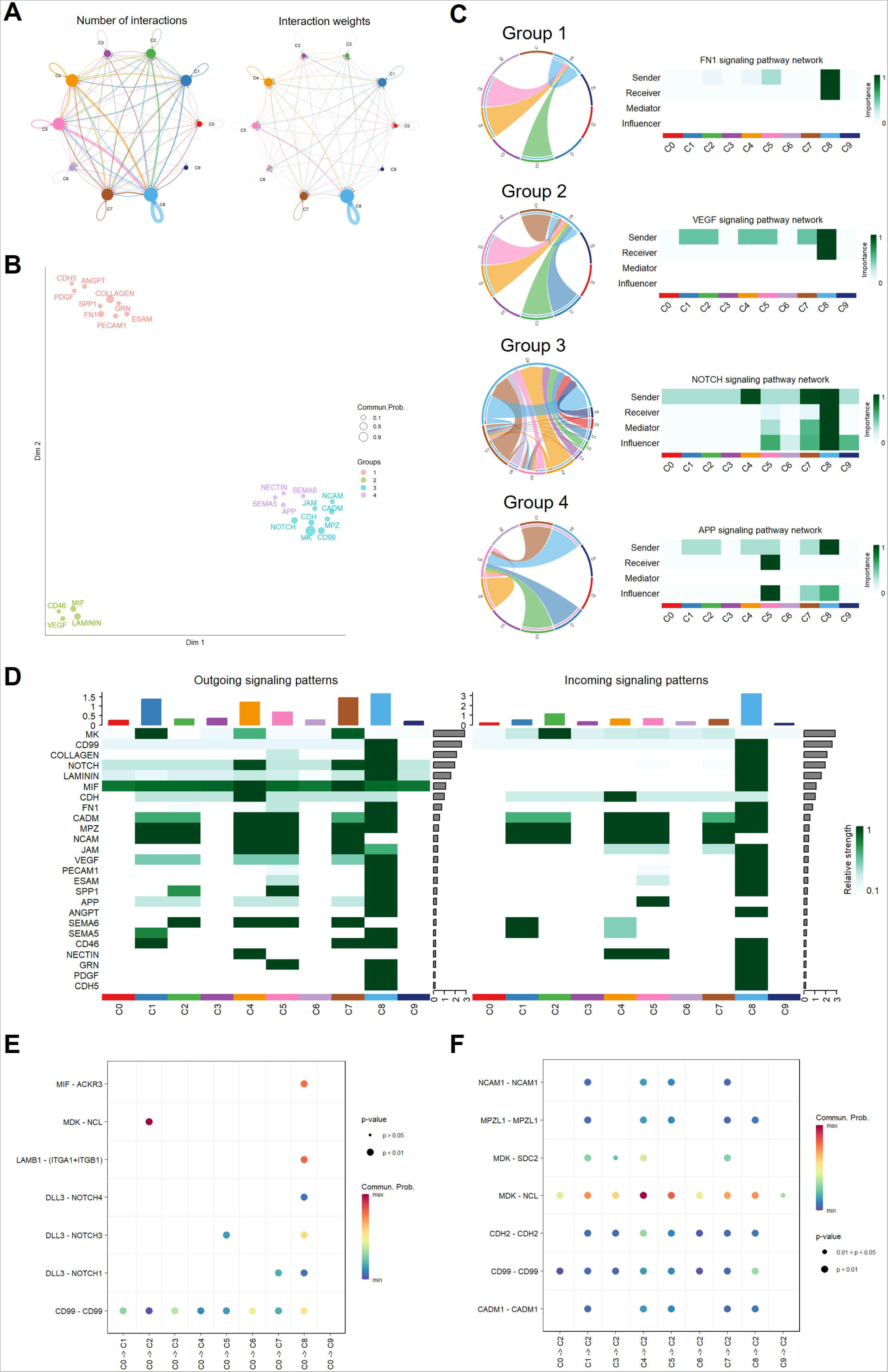
Cell-cell communications between RB subpopulations. A) Aggregate cell-cell communication network between RB clusters based on number of interactions (left) or total interaction strengths (right). B) Scatter plot showing grouping of signaling pathways based on cellular communication network similarity. C) Representative signaling pathways in Group 1 (*FN1*), Group 2 (*VEGF*), Group 3 (*NOTCH*), Group 4 (*APP*). Chord diagrams visualizing the cell-cell communication between RB clusters (left panel). Heatmap showing the network centrality scores which identified the signaling roles of RB clusters in the intercellular communication network (right panel). D) Heatmap depicting outgoing and incoming signaling patterns in the RB clusters. E-F) Dot plots showing significant ligand-receptor interactions for RB cluster interactions for E) the initial state C0 and F) terminal state C2.

We investigated the outgoing and incoming signaling patterns for individual RB clusters to dissect their roles in the communication network (Figure 8D). Interestingly, many of the identified pathways were involved in interactions with C8 cells, potentially highlighting a key role of glial cells in mediating cellular communication within the TME of RB. For example, *PECAM-1* is an integrin activator and an apoptosis suppressor that mediates both anchorage-independent growth and anoikis resistance in tumor cells ^74^. *PECAM-1* showed strong outgoing and incoming signals in C8 cells, indicating a self-self communication within C8 cells. In contrast, the *MK* pathway mediated communication from C1, C4, C7 to C2 . Also, C9 showed low activities in the overall outgoing and incoming signaling patterns, potentially reflecting a less prominent intercellular role of CAFs within the TME of RB (Figure 6D).

To dissect the signaling involved in RB progression, we analyzed the ligand-receptor pairs that drive outgoing signals for the initial RB state (C0 cells) and incoming signals for the terminal RB state (C2 cells, Figure 6E-F, Supplementary figure 6). We found that C0 communicates with C1-C8 via *CD99* signaling, which is a cell surface tumor marker involved in cell migration, tumor invasion and metastasis ^75^. Also, our results highlighted a number of additional pathways utilized by C0 to communicate with C8, including signaling via the *MIF-ACKR3* axis, *LAMB1-ITGA1/B1* axis and *DLL-NOTCH* axis. On the other hand, our results highlight a key role of *MDK-NCL* axis for the C2 terminal state to receive signals from other clusters. Midkine (*MDK*) is a neurotrophic factor known for maintaining cancer cell self-renewal and tumorigenic, while *NCL* is associated with early apoptotic cell recognition and immune activity suppression ^76,77^. Our result is consistent with a previous report that *MDK-NCL* represented a key pathway in conferring malignancy and suppressing TME immune activities ^78^.

## Discussion

Our current understanding of the molecular mechanisms underlying human RB progression largely came from scRNA-seq studies which lack spatial information. Here we report the first spatial transcriptomic profiling of primary RB tumors to create a spatial gene atlas, providing new insights into the spatial communication between RB subpopulations in the TME.

Our study provided a unique snapshot for the spatial pattern of a RB tumor sample from a 4-year-old patient presenting as a genetically unresolved case. Notably, we found a number of CNVs in the RB tumor, including the commonly found Chr1p and Chr16q amplification and Chr16p loss ^5^. Our results showed the tumor has high levels of *MYCN* expression and low *RB1* expression, suggesting a sporadic case of RB. Also, the majority of the tumor expressed *UBE2C*, a prognostic marker previously reported to activate malignant progression in RB ^11^. Consistent with previous scRNA-seq studies in RB ^5,11,12^, our results highlighted high intratumoral heterogeneity in RB. We identified 10 subpopulations representing different stages of RB with varying malignancy. Trajectory analysis unraveled the complex tumor cell trajectory linking RB cells at the initial state (eg C0) to the terminal state (C2). The gene cascades driving RB progression are notably associated with cancer-related functions, including cell proliferation, adhesion, tumor migration and invasion, and immune responses. We showed that the *JUND* and *SOX4* reglons are involved in the initial state, while *ETS1* and *CEBPD* regions are involved in the intermediate state, providing insights into the genetic regulation driving RB progression. Also, our data highlighted upregulation of cancer-related and prognosis marker genes in the terminal state C2, such as the immortal marker *MTRNR2L12* ^79^, *MALAT1* which is a cancer biomarker and associated in photoreceptor survival ^80,81^, *PLCG2* which is linked to metastasis ^82^ and the oncogene *YBX1* ^83^. Future implementation of multi-omics analysis in the same RB tumor, such as ATAC-seq or metabolomic, would complement the spatial transcriptomics findings and allow us to comprehensively profile RB.

Understanding the cell origin of RB is important to develop targeted therapy and minimize off-target issues. Early mouse studies suggest that RB can arise from amacrine cells^84^, horizontal cells^85^ or Müller glia cells ^86^. In contrast, recent studies point to cone precursors as the origin of RB in humans ^5,11,12,20^ . Consistent with this, we showed a dominant presence of cone precursors in RB tumor, as well as presence of highly proliferated cone precursors, CAFs and glial cells to form a complex TME. Our results also revealed a complex spatial communication network within the TME. For instance, C8 cells were identified as the key mediator of intercellular communication between RB subpopulations via a number of pathways, including *VEGF* and *NOTCH* signalings. Furthermore, our results supported a role of fibronectin (*FN1*) and IGF signaling (*IGFBP7*) in mediating the spatial communication of the terminal state C2 with its neighbors. Also, the *MDK-NCL* ligand-receptor pair was identified as a key pathway to mediate incoming signals in C2, consistent with the known role for midkine signaling driving cancer progression ^68^ . Together, our findings provide novel insights to understand the spatial communication landscape within the RB tumor.

There are limitations to this study. Previous studies have reported high variability among different RB tumors ^5^, partly due to the high number of oncogenic lesions that can lead to RB tumorigenesis. Here we have surveyed a single tumor sample and demonstrated the potential of using spatial transcriptomics to comprehensively analyze the spatial networks in the RB tumor. Future studies to profile more RB tumor samples could reveal new spatial communications in RB, however identification of a universal spatial communication network in RB might prove challenging given the high inter-tumor variability and would likely require a large scale of RB sampling. Also, this study is limited by the resolution of the Visium platform which did not reach true single cell level in human retinal cells. To address this, we utilized cell deconvolution using RB scRNA-seq dataset to resolve and improve the resolution of our spatial transcriptomic dataset. Alternative spatial transcriptomic systems could be applied to improve RB profiling in future research ^87^. For instance, imaging-based spatial systems such as MERSCOPE and Xenium offer spatial profiling at single cell level, but are greatly limited by the number of detected genes. In contrast, sequencing-based spatial systems such as Visium and GeoMx provide whole transcriptome analysis but can not reach single cell resolution. The recent system Stereo-seq has overcome this resolution hurdle, reaching a nanoscale resolution of 500nm. Future advances in spatial transcriptomic technology would facilitate large scale, high-resolution spatial profiling of RB tumors in a cost-effective manner.

In conclusion, we report the first spatial gene atlas for RB which represents a significant advancement in unraveling the intricacies of human RB progression. By mapping spatial intercellular networks involved in RB, this study provides a comprehensive insight into the complex spatial signaling that underlie pathological progression of RB tumor. Our study provides a compelling model to support the application of spatial transcriptomic to study the retina, and support future application of multi-omics approaches to comprehensively map the complex architecture of tumors.

## Methods

### Collection of patient samples

Collection of clinical donor samples was approved by the Human Research Ethics committee of the Royal Victorian Eye and Ear Hospital (HREC13/1151H) and the Royal Children’s Hospital Melbourne (HREC/78024/RCHM-2021), and carried out according to approved guidelines. Informed consent was obtained from all donors, and experiments conformed to the principles set out in the WMA Declaration of Helsinki. Sample freezing and OCT embedding followed the Visium protocol by 10X genomics. RB tumor sample was collected from patient enucleated eyes by the Children’s Cancer Centre Biobank (Royal Children’s Hospital and Murdoch Children’s Research Institute, HREC 33207) and immediately snap-frozen followed by embedding in OCT. The eye globes from three healthy donors with short post-mortem time were collected by the Lions Eye Donation Service (Royal Victorian Eye and Ear Hospital, Supplementary table 1). The eye globes were immediately simultaneously processed for OCT embedding and freezing to preserve the tissue structure.

### Spatial transcriptomic profiling using Visium

OCT-embedded retinal and RB samples were cryosectioned to 20 μm sections using a CryoStar NX70 (Thermo Fisher Scientific), and placed onto the capture areas of the Visium Spatial Tissue Optimisation and Visium Gene Expression slides following the manufacturer’s instructions. The sections were then fixed using methanol and stained with H&E following the Visium protocol. Images of the whole capture area were acquired using an Axio Imager M2 microscope (Zeiss) and stitched together with the tiles tool of ZEN blue edition (Zeiss). Tissue sections on the Visium Gene Expression slide were permeabilized for 24min to capture the mRNAs with probes on the gene expression spots. Reverse transcription mix was later added to the RB sections to synthesize double-stranded cDNA, followed by the preparation of the spatial transcriptome library using the 10x Genomics Visium Spatial Gene Expression Reagent Kits. The cDNA library was sequenced using NovaSeq with 150bp paired end sequencing, yielding ∼54-132M reads/section (Australian Genome Research Facility).

### Preprocessing and QC of Visium Profiles

Raw sequencing data in fastq format was aligned to the human genome reference GRCh38 using the 10x Genomics SpaceRanger pipeline (v2.0). This yielded gene expression matrices describing transcriptomic features according to the raw UMI counts of each spot, and position lists containing spatial features including barcodes and coordinates of each capture-spot on the 10x Visium slides. We further integrated and preprocessed the SpaceRanger outputs using STUtility (v0.1) and Seurat (v4.0) ^88,89^. Transcriptomic data from the sections were assigned to each spot according to the Visium barcodes and coordinates, and merged into a Seurat object for quality check using the STUtility pipeline. Valid spots were manually selected on the H&E images using the 10x Genomics Loupe Browser. Spots with greater than 30% mitochondrial content and expressing less than 20 unique genes (UMI) were excluded. After filtering, a Seurat object containing 2982 spots and 16398 unique genes was retained for RB, as well as 3761 spots and 19233 unique genes in the human retina.

### Image loading and processing

Brightfield images from the H&E staining were loaded into the Seurat object through *LoadImage*. For faster processing, scaled down versions (400 pixels wide) were automatically saved, while plots for publication were based on raw images with higher resolution. Background removal was achieved using a binary mask through *MaskImages*. Spots with transcriptomic profiles were then mapped to the masked images according to their corresponding coordinates. Then, *FeatureOverlay* computed the overlay of spatial features onto the H&E images, and *FeaturePlot* plotted specific features on the spot coordinates.

### Transcriptomic data normalization, dimensional reduction and clustering

Data was normalized by variance stabilizing transformation using *SCTransform* in Seurat (V4.0). To address sample section variations, the “section” was specified with the “vars.to.repress” option using *SCTransform* ^90^. Next, the normalized data was reduced into a lower dimensional space using principal components analysis (PCA). Our dataset was integrated from the transcriptome profiles of multiple sections, thus an additional harmony integration was applied for batch correction, aiming to yield an integrated embedding with clear transitions between clusters on the lower dimensional space. Harmony algorithm was applied by inputting the top 30 principal components (PCs) into *RunHarmony* within Seurat to smooth the transition between individual spots and integrate the discrepancies in the spatial distribution of clusters across sections ^91^. The Harmony-integrated spots were then embedded in a K-nearest neighbor (KNN) graph structure, with edges drawn between spots with similar feature patterns to partition the graph into highly interconnected clusters. *FindNeighbors* constructed a KNN graph based on the Euclidean distance in the Harmony-integrated PCA space, and calculated the neighborhood overlap between each cell and its local neighbors. Afterwards, louvain algorithm was performed for modularity optimization via *FindClusters* to iteratively group cells together at a resolution of 0.8 clustering granularity. Uniform manifold approximation and projection (UMAP) non-linear dimensional reduction algorithm was applied using *RunUMAP* in Seurat to place similar cells together in the low-dimensional space ^92^. Visualization and annotation of identified clusters on spot coordinates and H&E images were accomplished using the STUtility (V0.1) and Seurat (V4.0) packages. For 3D visualization, *Create3DStack* was used to create a z-stack of 2D point patterns, followed by interpolation of expression values across the point patterns and visualization using *FeaturePlot3D*.

### Cell type annotation and single cell deconvolution

Retinal markers were collated from previous publications to annotate the specific cell type of each cluster (Supplementary Table 1). Retinal markers were input as a list of gene expression patterns into *AddModuleScore* which calculate the average expression of each pattern for each cluster. The aggregated expression of randomly selected control gene sets was subtracted to generate a module score for each retinal cell type for each cluster, before using *VlnPlot* to visualize the module scores of each retinal cell type across all identified clusters.

Conditional autoregressive-based deconvolution (CARD) algorithm was used next to combine cell-type-specific transcriptomic profiles from reference single-cell RNA sequencing datasets with correlation in cell-type composition across spatial spots ^18^. The spatial location coordinates and single-cell RNA sequencing metadata were formatted into a data frame, while the spatial transcriptomic profiles and single-cell RNA sequencing count matrix were formatted into a sparse matrix. These datasets were input using *createCARDObject* in CARD (V1.0) to create a CARD object, followed by deconvoluting the spatial transcriptomic data based on an efficient optimization algorithm for constrained maximum likelihood estimation. The dominant cell type of each cluster was annotated by the top distributed cell type with a threshold of 0.5, as calculated by the average possibility of each cell type across all spots belonging to the same cluster which means the higher score, the likelier the proportion of corresponding cell types in a cluster ^93^. The threshold of minor/associated cell types was set to 0.1 to retain rare but significant cells ^94^.

Deconvolution results were visualized through the cell type proportions on spot coordinates using *CARD.visualize.prop*. The proportion of each cell type of interest in each cluster was manually calculated based on the average cell type probability across every spot in each cluster saved in the CARD object. The putative identity of each cell type in the reference scRNA-seq dataset was extracted according to the differentially expressed gene analysis pipeline. The distribution of identified gene signatures associated with specific cell types were dot-plotted across all 10 RB clusters to evaluate the deconvolution results.

### Cell cycle scoring

Cell cycle scoring calculated the average expression levels of canonical S and G2/M phase markers stored in Seurat across all spots captured on our RB sections. Cell cycle scores were plotted on spatial coordinates and the UMAP embedding according to Seurat vignettes. *RidgePlot* plotted the distribution of cell cycle markers of interest across all 10 clusters, and scores were compiled for a bar chart depicting the ratio of cells undergoing different cell cycle phases in each cluster.

### RNA velocity analysis and trajectory analysis

Spaceranger output files, genome annotation files and repeat masker files were transformed into loom files using *run10x* from the velocyto (V0.17.16) package ^22^ and loaded into python for RNA velocity analysis using scVelo (V0.2.5) ^95^. Reads in the loom files were filtered and normalized, before computing the average and uncentered variances among nearest neighbors in PCA space for velocity estimation using *scv.pp.filter_and_normalize* and *scv.pp.moments*. Velocities were analyzed by obtaining the stochastic model of transcriptional dynamics using *scv.tl.velocity*, and the combination of velocity vectors across genes estimated the future state of each spot. Cell transition probabilities were calculated via the cosine correlation between potential cell transitions, and the velocity vectors using *scv.tl.velocity_graph*. Velocities were projected onto UMAP embeddings as streamlines using the mean transition and transition probabilities via *scv.pl.velocity_embedding_stream*.

Next, CellRank (V1.5.1) estimated the initial and terminal states, as well as putative lineage drivers based on the Markov chain concept ^23^. A dynamical model of splicing kinetics was solved by iteratively estimating the parameters of reaction rates and latent cell-specific variables using *scv.tl.recover_dynamics*. Subsequently, directional information from velocity vectors was extracted and applied to compute the initial and terminal states from the set of macrostate via the stability criterion using *cr.tl.initial_states* and *cr.tl.terminal_states*. Lineage driver genes were identified using *cr.tl.lineage_drivers* from CellRank.

The identification of initial and terminal states in our RB sections provided information about root cells and diffusion lineages, necessary for computing velocity pseudotime using *scv.pl.velocity_pseudotime*, which was plotted on the spatial coordinates and UMAP embedding. The identification of top driven genes and the velocity pseudotime along cell fate trajectories later helped define the fate map and molecular mechanism driving specific RB lineages. Lineage drivers were sorted by their confidence along the fate trajectories using *cr.pl.heatmap*. Finally, the topology of connectivity between each cluster and trajectory inference were summarized by a PAGA graph superimposed onto the UMAP embedding using *scv.tl.paga* ^30^.

### CNV analysis

CNV was calculated using the inferCNV package (v1.19.1) in R, which identified gains or deletions of whole chromosomes or large chromosomal segments ^96^. The spatial transcriptomic profiles of control human retina served as the reference normal cells to explore the expression intensity of genes in RB clusters across positions of the genome compared to the reference set. *infercnv:run* was used for the estimation, with parameters set as denoise = T, HMM = T, and a cutoff of 0.1. The specific CNV scores for each cell were calculated as mean squares of CNV values across the genome ^16^.

### DE gene analysis and Gene ontology (GO) analysis

DE genes that define clusters were identified using *FindAllMarkers* which performed non-parametric Wilcoxon Rank Sum tests between each cluster to all other spots. The log fold change threshold was set to 0.25 to only consider unique genes present in at least 25% of spots in each cluster. *DoHeatmap* from the Seurat package then plotted the expression heatmap of the top 10 markers. Biological functions of the identified DE genes were further probed using the clusterProfiler package (V4.6) ^97^. Using clusterProfiler vignettes, the input data was constructed using a list of marker gene names and corresponding log2 fold change values, which was then annotated by GO terms based on Biological Processes using *gseGO*. Finally, the enriched GO Biological Processes terms were visualized using dot plots and category netplots.

### SCENIC analysis

Transcriptomic profiles were first filtered using *runGenie3* of the GENIE3 package for potential targets for each TF based on co-expression ^41^. Next, potential direct- and indirect-binding targets (regulons) were selected from the co-expression network through cis-regulatory motif analysis using *runSCENIC_2_createRegulons* of the RcisTarget package. The regulon events were then scored in individual cells using *runSCENIC_3_scoreCells* of the AUCell package. Finally, the gene regulatory network activity was combined with clusterings using *regulonAUC* to identify the top regulons in cell transitions, and explore the association of cell states with specific regulons.

### Neighborhood expression analysis

*RegionNeighbours* from the STUtility package was used to identify and select spots neighboring to our clusters of interest. Transcriptomic profiles at the neighboring borders were later extracted for DE gene analysis, before performing GO terms enrichment analysis following the workflow described above.

### Cell-cell communication analysis

The CellChat (V1.6.1) pipeline was used to model the probability of cell-cell communication ^48^. First, the gene expression data and cell meta information were used to create a CellChat object using *createCellChat*. Next, we utilized *CellChatDB.human*, which is a literature-supported database of 1939 published molecular interactions, as the reference ligand-receptor interaction database. Cell state-specific communications were identified using *identifyOverExpressedGenes* and *identifyOverExpressedInteractions* by assessing ligand or receptor overexpression in each cluster, followed by *computeCommunProb* to inferred cellular communication networks, with the default calculation set to ‘trimean’ for more accurate estimations, and *computeCommunProbPathway* computed communication probabilities at the pathway level. Finally, the aggregated cell-cell communication networks were computed based on the communication probabilities across all estimated ligand-receptor pairs and signaling pathways via *aggregateNet*.

Chord diagrams and network centrality scores measuring out-degree, in-degree, flow betweenness, and information centrality were generated using *netVisual_aggregate* and *netAnalysis_computeCentrality*. This identified dominant senders, receivers, mediators, and influencers of the intercellular communications for visualization in heatmaps by *netAnalysis_signalingRole_network* for each signaling pathway across all 10 clusters. CellChat facilitated the interpretation of complex networks by predicting key incoming and outgoing signals for each cluster along with coordinated responses among different clusters. This was achieved, via *netAnalysis_signalingRole_heatmap* that used pattern recognition to group pathways based on similarity measures, and *computeNetSimilarity* that performed manifold learning from a functional perspective. Finally, significant ligand-receptor pairs contributing to incoming signals to the terminal state, and outcoming signals from the initial state were visualized using *netVisual_bubble*.

## Acknowledgement

We would like to thank the Lions Eye Donation Services and the Royal Victorian Eye and Ear Hospital for the collection of donor eye samples. This research was funded by the Bayer Global Ophthalmology Awards Program (SH, RCBW, AHW, SS). RCBW is supported by the University of Melbourne, and the Centre for Eye Research Australia, National Health and Medical Research Council (GCT1184076) and Medical Future Research Fund (MRF2024365). DU is supported by the Melbourne Research Scholarship. The Centre for Eye Research Australia and St Vincent Institute of Medical Research receive operational infrastructure support from the Victorian Government. AGRF is supported by the Australian Government National Collaborative Research Infrastructure Strategy through Bioplatforms Australia. Tumor samples and coded data were supplied by the Children’s Cancer Centre Biobank at the Murdoch Children’s Research Institute and The Royal Children’s Hospital (mcri.edu.au/research/projects/childrens-cancer-centre-biobank). Establishment and running of the Children’s Cancer Centre Biobank is made possible through generous support by Cancer In Kids @ RCH (www.cika.org.au), The Royal Children’s Hospital Foundation and the Murdoch Children’s Research Institute.

## Data availability

The sequencing data in this study are available in the Human Cell Atlas data portal.

## Code availability

The code for data analysis is available at Github (https://github.com/wongcb/Spatial-transcriptomic-retinoblastoma).

## References

1. Kivelä, T. The epidemiological challenge of the most frequent eye cancer: retinoblastoma, an issue of birth and death. The British journal of ophthalmology vol. 93 1129–1131 (2009).

2. Rushlow, D. E. et al. Characterisation of retinoblastomas without RB1 mutations: genomic, gene expression, and clinical studies. Lancet Oncol. 14, 327–334 (2013).

3. Afshar, A. R. et al. Next-Generation Sequencing of Retinoblastoma Identifies Pathogenic Alterations beyond RB1 Inactivation That Correlate with Aggressive Histopathologic Features. Ophthalmology 127, 804–813 (2020).

4. Zhang, J. et al. A novel retinoblastoma therapy from genomic and epigenetic analyses. Nature 481, 329–334 (2012).

5. Wu, C. et al. Single-cell characterization of malignant phenotypes and microenvironment alteration in retinoblastoma. Cell Death Dis. 13, 438 (2022).

6. Singh, H. P. et al. Developmental stage-specific proliferation and retinoblastoma genesis in RB-deficient human but not mouse cone precursors. Proc. Natl. Acad. Sci. U. S. A. 115, E9391–E9400 (2018).

7. Macpherson, D. Insights from mouse models into human retinoblastoma. Cell Div. 3, 9 (2008).

8. Aldiri, I. et al. The Dynamic Epigenetic Landscape of the Retina During Development, Reprogramming, and Tumorigenesis. Neuron 94, 550–568.e10 (2017).

9. Liu, H. et al. Human embryonic stem cell-derived organoid retinoblastoma reveals a cancerous origin. Proc. Natl. Acad. Sci. U. S. A. 117, 33628–33638 (2020).

10. Norrie, J. L. et al. Retinoblastoma from human stem cell-derived retinal organoids. Nat. Commun. 12, 4535 (2021).

11. Yang, J. et al. Single-cell transcriptome profiling reveals intratumoural heterogeneity and malignant progression in retinoblastoma. Cell Death Dis. 12, 1100 (2021).

12. Collin, J. et al. Dissecting the Transcriptional and Chromatin Accessibility Heterogeneity of Proliferating Cone Precursors in Human Retinoblastoma Tumors by Single Cell Sequencing-Opening Pathways to New Therapeutic Strategies? Invest. Ophthalmol. Vis. Sci. 62, 18 (2021).

13. Hunter, M. V., Moncada, R., Weiss, J. M., Yanai, I. & White, R. M. Spatially resolved transcriptomics reveals the architecture of the tumor-microenvironment interface. Nat. Commun. 12, 6278 (2021).

14. Arora, R. et al. Spatial transcriptomics reveals distinct and conserved tumor core and edge architectures that predict survival and targeted therapy response. Nat. Commun. 14, 5029 (2023).

15. Patel, A. P. et al. Single-cell RNA-seq highlights intratumoral heterogeneity in primary glioblastoma. Science 344, 1396–1401 (2014).

16. Sun, K. et al. scRNA-seq of gastric tumor shows complex intercellular interaction with an alternative T cell exhaustion trajectory. Nat. Commun. 13, 4943 (2022).

17. Qi, D.-L. & Cobrinik, D. MDM2 but not MDM4 promotes retinoblastoma cell proliferation through p53-independent regulation of MYCN translation. Oncogene vol. 36 1760–1769 Preprint at 10.1038/onc.2016.350 (2017).

18. Ma, Y. & Zhou, X. Spatially informed cell-type deconvolution for spatial transcriptomics. Nat. Biotechnol. 40, 1349–1359 (2022).

19. Li, H. et al. A comprehensive benchmarking with practical guidelines for cellular deconvolution of spatial transcriptomics. Nat. Commun. 14, 1548 (2023).

20. Xu, X. L. et al. Rb suppresses human cone-precursor-derived retinoblastoma tumours. Nature 514, 385–388 (2014).

21. Hanahan, D. Hallmarks of Cancer: New Dimensions. Cancer Discov. 12, 31–46 (2022).

22. La Manno, G. et al. RNA velocity of single cells. Nature 560, 494–498 (2018).

23. Lange, M. et al. CellRank for directed single-cell fate mapping. Nat. Methods 19, 159–170 (2022).

24. Setty, M. et al. Characterization of cell fate probabilities in single-cell data with Palantir. Nat. Biotechnol. 37, 451–460 (2019).

25. Hartford, C. C. R. et al. Context-Dependent Function of Long Noncoding RNA in Transcriptome Regulation during p53 Activation. Mol. Cell. Biol. 42, e0028922 (2022).

26. Geng, L., Wang, Z. & Tian, Y. Down-regulation of ZNF252P-AS1 alleviates ovarian cancer progression by binding miR-324-3p to downregulate LY6K. J. Ovarian Res. 15, 1 (2022).

27. Lu, Q., Ding, Y., Li, Y. & Lu, Q. 5-HT receptor agonist Valerenic Acid enhances the innate immunity signal and suppresses glioblastoma cell growth and invasion. Int. J. Biol. Sci. 16, 2104–2115 (2020).

28. Zhang, W., Li, L., Bian, P.-P., Luo, Q.-P. & Xiong, Z.-T. PLEKHA4 Is a Prognostic Biomarker and Correlated with Immune Infiltrates in Glioma. Biomed Res. Int. 2023, 4504474 (2023).

29. Qian, M. et al. PLEKHG5 is a novel prognostic biomarker in glioma patients. Int. J. Clin. Oncol. 24, 1350–1358 (2019).

30. Wolf, F. A. et al. PAGA: graph abstraction reconciles clustering with trajectory inference through a topology preserving map of single cells. Genome Biol. 20, 59 (2019).

31. Ko, M. J. et al. RPL17 Promotes Colorectal Cancer Proliferation and Stemness through ERK and NEK2/β-catenin Signaling Pathways. J. Cancer 13, 2570–2583 (2022).

32. Dong, M. et al. H2AFZ Is a Prognostic Biomarker Correlated to TP53 Mutation and Immune Infiltration in Hepatocellular Carcinoma. Front. Oncol. 11, 701736 (2021).

33. Mascarenhas, J. B. et al. EVL is a novel focal adhesion protein involved in the regulation of cytoskeletal dynamics and vascular permeability. Pulm. Circ. 11, 20458940211049002 (2021).

34. Stingl, K. et al. CDHR1 mutations in retinal dystrophies. Sci. Rep. 7, 6992 (2017).

35. Ogawa, M. et al. ATP generation in a host cell in early-phase infection is increased by upregulation of cytochrome c oxidase activity via the p2 peptide from human immunodeficiency virus type 1 Gag. Retrovirology 12, 97 (2015).

36. McKenzie, M. & Duchen, M. R. Impaired Cellular Bioenergetics Causes Mitochondrial Calcium Handling Defects in MT-ND5 Mutant Cybrids. PLoS One 11, e0154371 (2016).

37. Ishii, K. et al. Characteristics and clustering of human ribosomal protein genes. BMC Genomics 7, 37 (2006).

38. Luo, G.-P., Ni, B., Yang, X. & Wu, Y.-Z. von Willebrand factor: more than a regulator of hemostasis and thrombosis. Acta Haematol. 128, 158–169 (2012).

39. Wang, H. et al. FN1 is a prognostic biomarker and correlated with immune infiltrates in gastric cancers. Front. Oncol. 12, 918719 (2022).

40. Necula, L. et al. Collagen Family as Promising Biomarkers and Therapeutic Targets in Cancer. Int. J. Mol. Sci. 23, (2022).

41. Van de Sande, B. et al. A scalable SCENIC workflow for single-cell gene regulatory network analysis. Nat. Protoc. 15, 2247–2276 (2020).

42. Elliott, B. et al. Essential role of JunD in cell proliferation is mediated via MYC signaling in prostate cancer cells. Cancer Lett. 448, 155–167 (2019).

43. Dittmer, J. The role of the transcription factor Ets1 in carcinoma. Semin. Cancer Biol. 35, 20–38 (2015).

44. Ko, C.-Y., Chang, W.-C. & Wang, J.-M. Biological roles of CCAAT/Enhancer-binding protein delta during inflammation. J. Biomed. Sci. 22, 6 (2015).

45. Liu, J. et al. SOX4 maintains the stemness of cancer cells via transcriptionally enhancing HDAC1 revealed by comparative proteomics study. Cell Biosci. 11, 23 (2021).

46. Kariminejad, A. et al. Retinitis pigmentosa, cutis laxa, and pseudoxanthoma elasticum-like skin manifestations associated with GGCX mutations. J. Invest. Dermatol. 134, 2331–2338 (2014).

47. Miura, T. et al. Single-Cell RNA-Seq Reveals LRRC75A-Expressing Cell Population Involved in VEGF Secretion of Multipotent Mesenchymal Stromal/Stem Cells Under Ischemia. Stem Cells Transl. Med. 12, 379–390 (2023).

48. Jin, S. et al. Inference and analysis of cell-cell communication using CellChat. Nat. Commun. 12, 1088 (2021).

49. Zhang, Y., Kontos, C. D., Annex, B. H. & Popel, A. S. Angiopoietin-Tie Signaling Pathway in Endothelial Cells: A Computational Model. iScience 20, 497–511 (2019).

50. Farooqi, A. A. & Siddik, Z. H. Platelet-derived growth factor (PDGF) signalling in cancer: rapidly emerging signalling landscape. Cell Biochem. Funct. 33, 257–265 (2015).

51. Inokuchi, M. et al. Cadherin 5 Is a Significant Risk Factor for Hematogenous Recurrence and a Prognostic Factor in Locally Advanced Gastric Cancer. Anticancer Res. 37, 6807– 6813 (2017).

52. Li, B. et al. Fibronectin 1 promotes melanoma proliferation and metastasis by inhibiting apoptosis and regulating EMT. Onco. Targets. Ther. 12, 3207–3221 (2019).

53. Abraham, V. et al. Involvement of TIMP-1 in PECAM-1-mediated tumor dissemination. Int. J. Oncol. 53, 488–502 (2018).

54. Ueda, T. et al. Endothelial Cell-Selective Adhesion Molecule Contributes to the Development of Definitive Hematopoiesis in the Fetal Liver. Stem Cell Reports 13, 992– 1005 (2019).

55. Arechavaleta-Velasco, F., Perez-Juarez, C. E., Gerton, G. L. & Diaz-Cueto, L. Progranulin and its biological effects in cancer. Med. Oncol. 34, 194 (2017).

56. Liu, Y. et al. A pan-cancer analysis of the oncogenic role of secreted phosphoprotein 1 (SPP1) in human cancers. Ann Transl Med 10, 279 (2022).

57. Rømer, A. M. A., Thorseth, M.-L. & Madsen, D. H. Immune Modulatory Properties of Collagen in Cancer. Front. Immunol. 12, 791453 (2021).

58. Yang, X. et al. VEGF-B promotes cancer metastasis through a VEGF-A-independent mechanism and serves as a marker of poor prognosis for cancer patients. Proc. Natl. Acad. Sci. U. S. A. 112, E2900–9 (2015).

59. Elvington, M., Liszewski, M. K. & Atkinson, J. P. CD46 and Oncologic Interactions: Friendly Fire against Cancer. Antibodies (Basel) 9, (2020).

60. Givant-Horwitz, V., Davidson, B. & Reich, R. Laminin-induced signaling in tumor cells. Cancer Lett. 223, 1–10 (2005).

61. Guda, M. R. et al. Pleiotropic role of macrophage migration inhibitory factor in cancer. Am. J. Cancer Res. 9, 2760–2773 (2019).

62. Anusewicz, D., Orzechowska, M. & Bednarek, A. K. Notch Signaling Pathway in Cancer-Review with Bioinformatic Analysis. Cancers 13, (2021).

63. Manara, M. C., Pasello, M. & Scotlandi, K. CD99: A Cell Surface Protein with an Oncojanus Role in Tumors. Genes 9, (2018).

64. Guan, F., Wang, X. & He, F. Promotion of cell migration by neural cell adhesion molecule (NCAM) is enhanced by PSA in a polysialyltransferase-specific manner. PLoS One 10, e0124237 (2015).

65. Lauko, A., Mu, Z., Gutmann, D. H., Naik, U. P. & Lathia, J. D. Junctional Adhesion Molecules in Cancer: A Paradigm for the Diverse Functions of Cell-Cell Interactions in Tumor Progression. Cancer Res. 80, 4878–4885 (2020).

66. Li, H., Gao, J. & Zhang, S. Functional and Clinical Characteristics of Cell Adhesion Molecule CADM1 in Cancer. Front Cell Dev Biol 9, 714298 (2021).

67. Luo, S., Lin, R., Liao, X., Li, D. & Qin, Y. Identification and verification of the molecular mechanisms and prognostic values of the cadherin gene family in gastric cancer. Sci. Rep. 11, 23674 (2021).

68. Filippou, P. S., Karagiannis, G. S. & Constantinidou, A. Midkine (MDK) growth factor: a key player in cancer progression and a promising therapeutic target. Oncogene 39, 2040–2054 (2020).

69. Chang, E. H. et al. Aminosalicylic acid reduces ER stress and Schwann cell death induced by MPZ mutations. Int. J. Mol. Med. 44, 125–134 (2019).

70. Purohit, A., Sadanandam, A., Myneni, P. & Singh, R. K. Semaphorin 5A mediated cellular navigation: connecting nervous system and cancer. Biochim. Biophys. Acta 1846, 485–493 (2014).

71. Wang, B. et al. The Amyloid Precursor Protein Is a Conserved Receptor for Slit to Mediate Axon Guidance. eNeuro 4, (2017).

72. Alto, L. T. & Terman, J. R. Semaphorins and their Signaling Mechanisms. Methods Mol. Biol. 1493, 1–25 (2017).

73. Maurel, P. et al. Nectin-like proteins mediate axon Schwann cell interactions along the internode and are essential for myelination. J. Cell Biol. 178, 861–874 (2007).

74. Zhang, X., Xu, L.-H. & Yu, Q. Cell aggregation induces phosphorylation of PECAM-1 and Pyk2 and promotes tumor cell anchorage-independent growth. Mol. Cancer 9, 7 (2010).

75. Pasello, M., Manara, M. C. & Scotlandi, K. CD99 at the crossroads of physiology and pathology. J. Cell Commun. Signal. 12, 55–68 (2018).

76. López-Valero, I. et al. Midkine signaling maintains the self-renewal and tumorigenic capacity of glioma initiating cells. Theranostics 10, 5120–5136 (2020).

77. Sorrelle, N., Dominguez, A. T. A. & Brekken, R. A. From top to bottom: midkine and pleiotrophin as emerging players in immune regulation. J. Leukoc. Biol. 102, 277–286 (2017).

78. Yu, X. et al. Integrating single-cell RNA-seq and spatial transcriptomics reveals MDK-NCL dependent immunosuppressive environment in endometrial carcinoma. Front. Immunol. 14, 1145300 (2023).

79. Song, L. et al. Cellular heterogeneity landscape in laryngeal squamous cell carcinoma. Int. J. Cancer 147, 2879–2890 (2020).

80. Li, Z.-X. et al. MALAT1: a potential biomarker in cancer. Cancer Manag. Res. 10, 6757– 6768 (2018).

81. Lukowski, S. W. et al. A single-cell transcriptome atlas of the adult human retina. EMBO J. 38, e100811 (2019).

82. Chan, J. M. et al. Signatures of plasticity, metastasis, and immunosuppression in an atlas of human small cell lung cancer. Cancer Cell 39, 1479–1496.e18 (2021).

83. Prabhu, L. et al. Role of post-translational modification of the Y box binding protein 1 in human cancers. Genes Dis 2, 240–246 (2015).

84. Dyer, M. A. & Bremner, R. The search for the retinoblastoma cell of origin. Nat. Rev. Cancer 5, 91–101 (2005).

85. Ajioka, I. et al. Differentiated horizontal interneurons clonally expand to form metastatic retinoblastoma in mice. Cell 131, 378–390 (2007).

86. Pajovic, S. et al. The TAg-RB murine retinoblastoma cell of origin has immunohistochemical features of differentiated Muller glia with progenitor properties. Invest. Ophthalmol. Vis. Sci. 52, 7618–7624 (2011).

87. Wang, Y. et al. Spatial transcriptomics: Technologies, applications and experimental considerations. Genomics 115, 110671 (2023).

88. Hao, Y. et al. Integrated analysis of multimodal single-cell data. Cell 184, 3573–3587.e29 (2021).

89. Bergenstråhle, J., Larsson, L. & Lundeberg, J. Seamless integration of image and molecular analysis for spatial transcriptomics workflows. BMC Genomics 21, 482 (2020).

90. Hafemeister, C. & Satija, R. Normalization and variance stabilization of single-cell RNA-seq data using regularized negative binomial regression. Genome Biol. 20, 296 (2019).

91. Korsunsky, I. et al. Fast, sensitive and accurate integration of single-cell data with Harmony. Nat. Methods 16, 1289–1296 (2019).

92. McInnes, L., Healy, J., Saul, N. & Großberger, L. UMAP: Uniform Manifold Approximation and Projection. J. Open Source Softw. 3, 861 (2018).

93. Xu, Z. et al. Integrative analysis of spatial transcriptome with single-cell transcriptome and single-cell epigenome in mouse lungs after immunization. iScience 25, 104900 (2022).

94. Coutant, A. et al. Spatial Transcriptomics Reveal Pitfalls and Opportunities for the Detection of Rare High-Plasticity Breast Cancer Subtypes. Lab. Invest. 103, 100258 (2023).

95. Bergen, V., Lange, M., Peidli, S., Wolf, F. A. & Theis, F. J. Generalizing RNA velocity to transient cell states through dynamical modeling. Nat. Biotechnol. 38, 1408–1414 (2020).

96. Tirosh, I. et al. Dissecting the multicellular ecosystem of metastatic melanoma by single-cell RNA-seq. Science 352, 189–196 (2016).

97. Yu, G., Wang, L.-G., Han, Y. & He, Q.-Y. clusterProfiler: an R package for comparing biological themes among gene clusters. OMICS 16, 284–287 (2012).

