## Supplementary figure 1-6, supplementary table 1-3 for "Spatial transcriptomic profiling of human retinoblastoma"

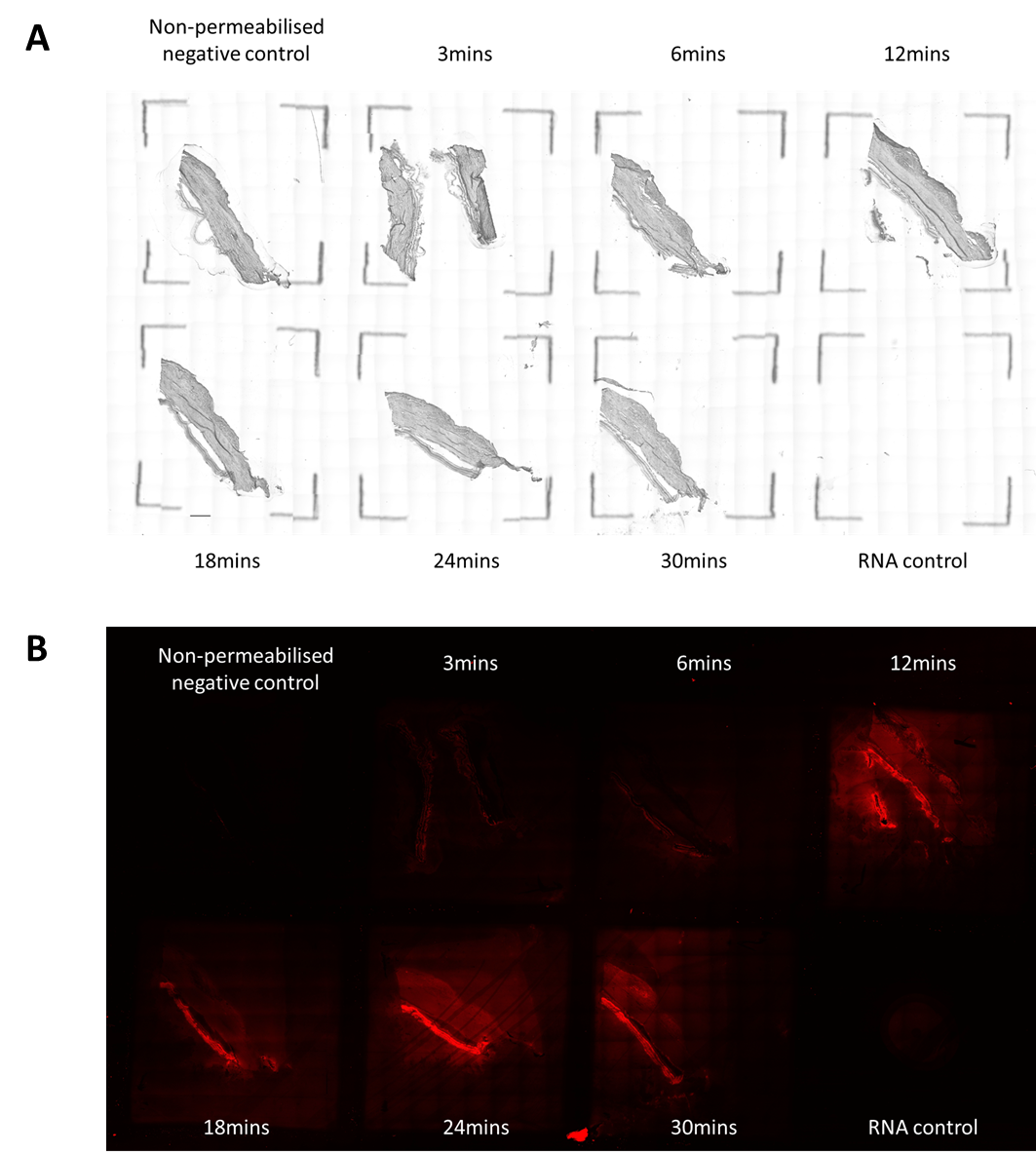

**Supplementary figure 1:** Tissue optimisation assay on donor retina. Control retina were mounted onto the Visium Tissue Optimisation slide and stained with A) hematoxylin and eosin. The tissues were then permeabilized at 0, 3, 6, 12, 18, 24 and 30 minutes and the amount of captured RNA from the tissue sample was determined by B) fluorescently labeling the RNA with TRITC probe.

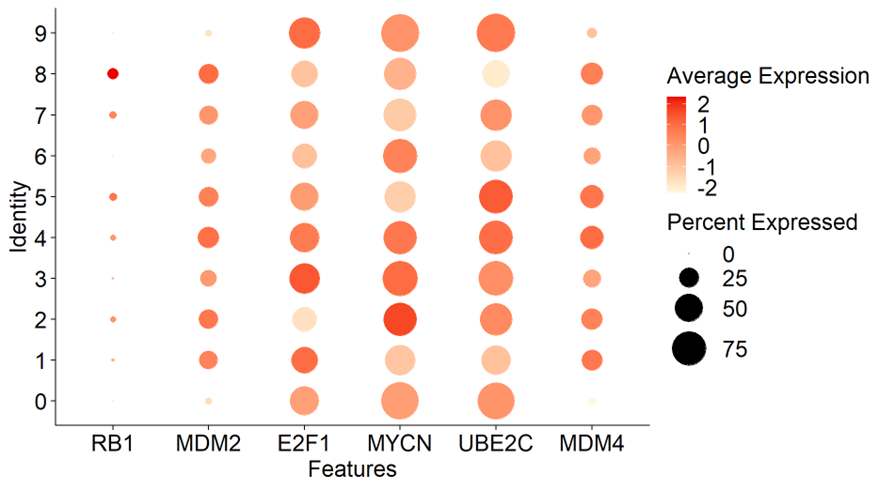

**Supplementary Figure 2:** Dot plot showing the expression levels of cancer-related genes *RB1, MDM2, E2F1, MYCN, UBE2C, MDM4* in RB.

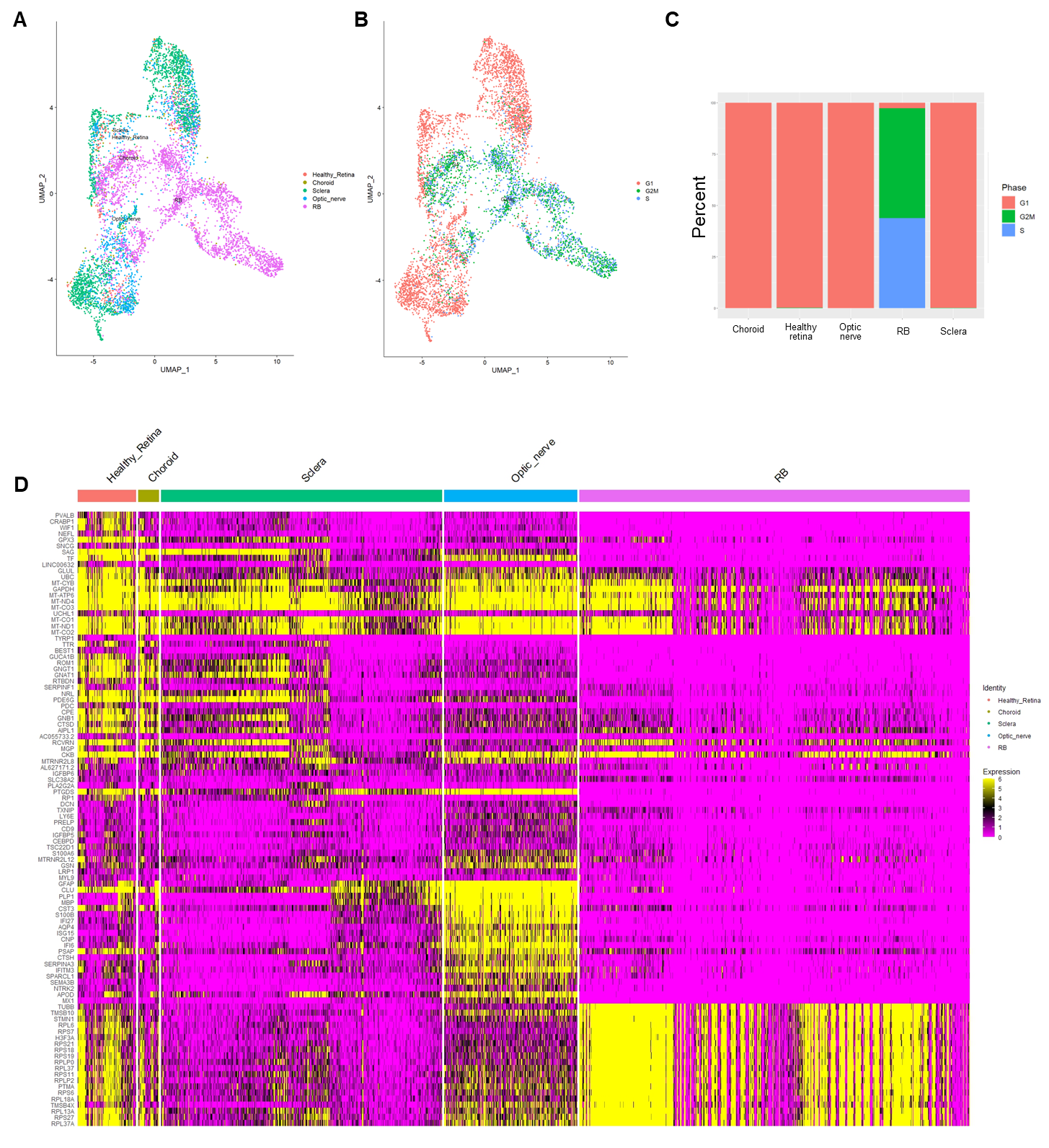

**Supplementary figure 3:** UMAP plot of combined dataset of A) control human retina (healthy retina, choroid, sclera, optic nerve) and RB tumor, as well as their B) cell cycle stages. C) Cell cycle analysis of human retina and RB. D) Heatmap showing expression levels of top DE genes in control retinal cell types and RB.

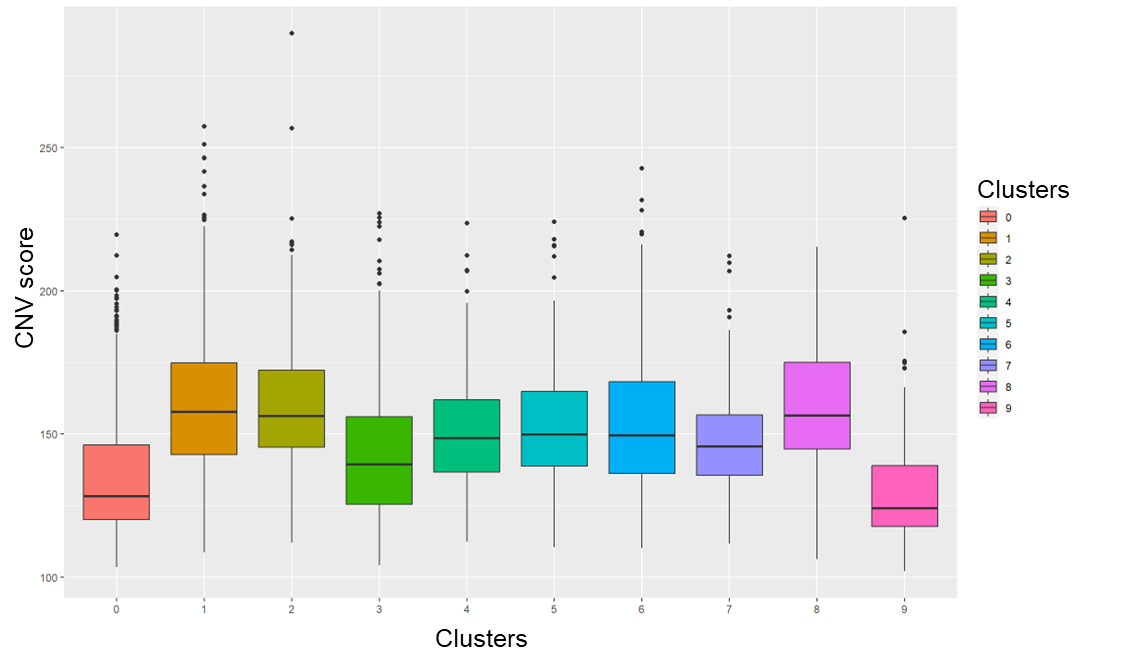

**Supplementary figure 4:** CNV score calculated by the mean squares of CNV values across the genome for RB clusters.

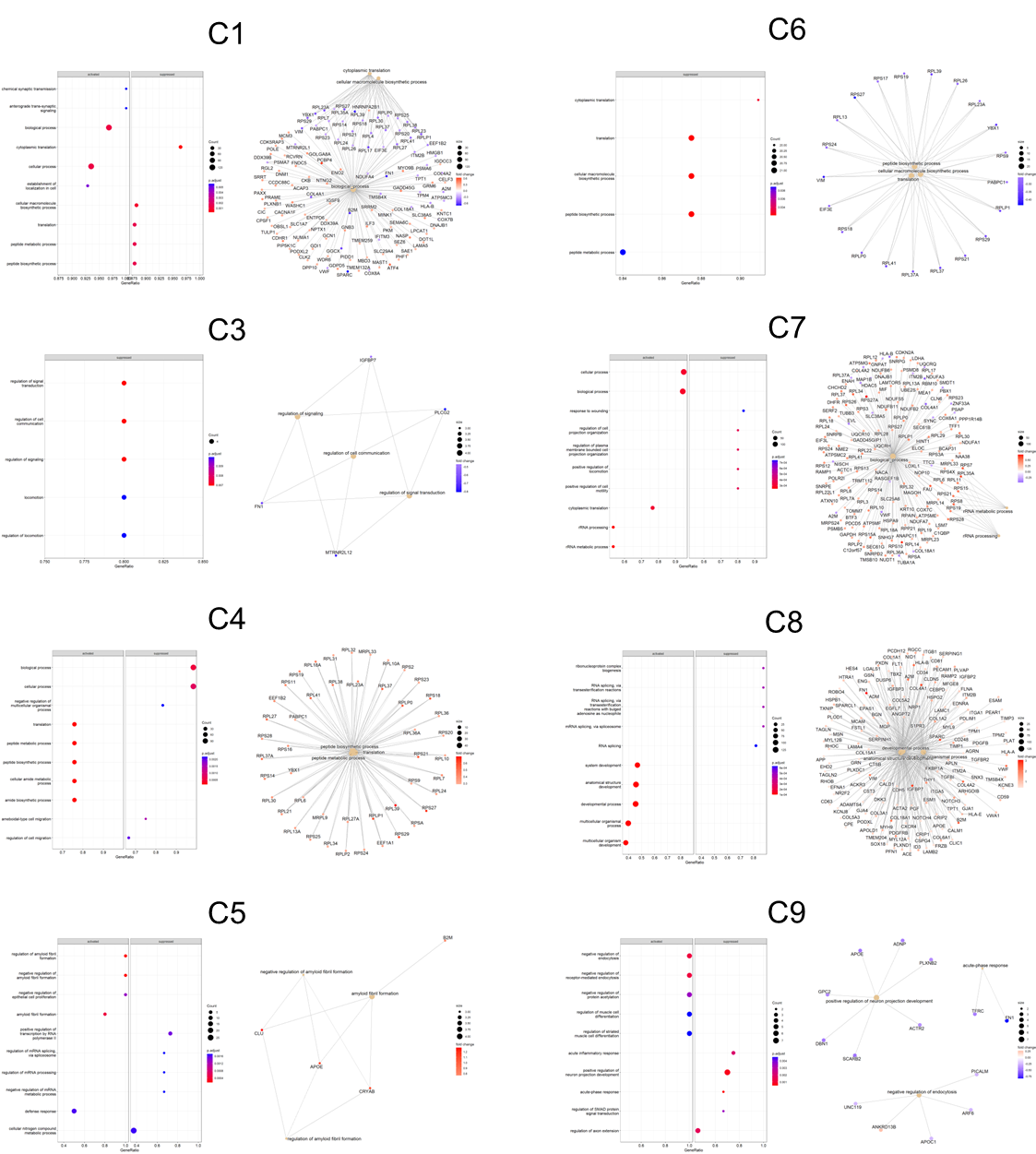

**Supplementary Figure 5.** Top GO Biological Process terms enriched in RB clusters as shown by dot plot (left panel) and category net plot (right panel).

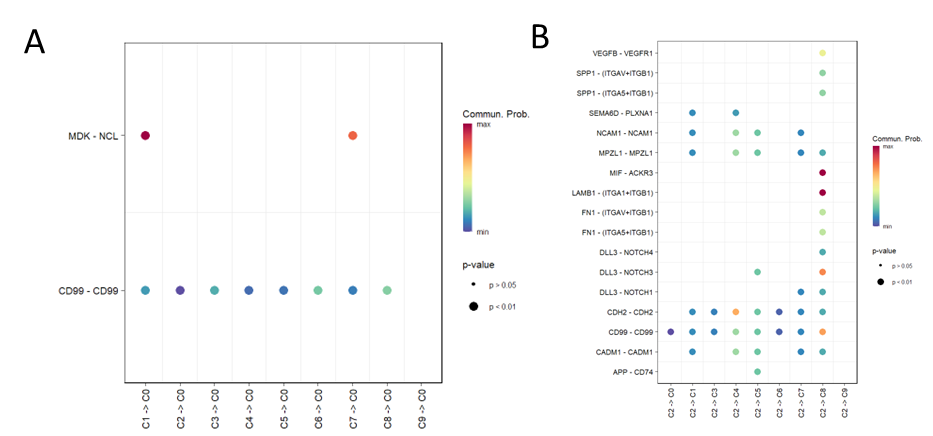

**Supplementary figure 6:** A) Incoming ligand-receptor interactions for the initial state C0 and B) outgoing ligand-receptor interactions for the terminal state C2.

**Supplementary table 1:** Information for donor samples used in this study.

| **Sample** | **Age (yrs)** | **Sex** | **Retrieval time (hrs)** | **Ocular history** | **Process** |
| --- | --- | --- | --- | --- | --- |
| RB | 4 | Male | 0 | RB - genetically unresolved | Visium |
| Control 1 | 47 | Male | 4 | Nil | Visium |
| Control 2 | 63 | Male | 10 | Cataracts, phakic eyes | Permeabilization optimization and Visium |
| Control 3 | 82 | Male | 8 | Bilateral intraocular lens | Visium |

**Supplementary Table 2. Marker genes for retinal cell type annotation**

| **Retinal cell types** | **Marker genes** |
| --- | --- |
| Retinal progenitor cell | *SOX2, HES1, MKI67, HES5, FZD5, PAX6* |
| Pigment epithelium cell | *SERPINF1, MITF, BEST1, TTR* |
| Rod cell | *RHO, PDE6A, CNGA1, NRL, GNAT1, GNB1, SAG, ELOVL4, PDE6B, GNGT1* |
| Cone cell | *ARR3, GNGT2, PDE6H, GUCA1C, GNAT2, RXRG, THRB, PDC, GNB3, CRX* |
| Horizontal cell | *ONECUT1, ONECUT2, TFAP2B* |
| Amacrine cell | *GAD1, CALB1, NRXN2, TFAP2A, PROX1, GAD2* |
| Retinal ganglion cell | *POU4F2, GAP43, NEFL, SNCG, ATOH7, EBF3, THY1, NRN1* |
| Bipolar cell | *GRM6, VSX2* |
| Cone precursor cell (CP) | *CRX, RXRG, THRB* |
| Highly-proliferated cone precursor (HP-CP) | *MKI67, TOP2A, KIF14, CRX, RXRG, THRB* |
| Glial cell | *CD68, HLA-DPA1, HLA-DPB1, CLU, APOE, GLUL* |
| Cancer-associated fibroblast (CAF) | *ACTA2, VIM, FGF9* |

**Supplementary Table 3. Identified regulons in RB clusters**

| **Regulon set** | **High-confident TF** | **Low-confident TF** | **Enriched Genes** |
| --- | --- | --- | --- |
| *JUND* | *BACH1; BACH2; MAF; MAFB; MAFF; MAFG; MAFK; NFE2; NFE2L1; NFE2L2; NFE2L3; PURA* | *EP300; FOS; FOSB; FOSL1; JUN; JUNB; JUND; KLF4; NR3C1; PAX4; RARA; RARB; RARG; RXRA; RXRB; RXRG; SP1; SP2; SP3; SP4; SREBF1; SREBF2; THRA; THRB; ZBTB14* | *ATRN, CCDC120, CD44, CRY2, DAG1, DERL3, DHRS11, DLG4, DZIP1L, EIF5AL1, FKTN, FOXP1, FRMD6, GPR162, HAUS7, KCNAB2, L1CAM, LIN28B, LMNA, LRRC4, MEIS3, NR4A1, P4HA2, PLEKHG5, POLR1E, REEP6, SEMA6C, SFXN3, SLC12A4, SNN, TIAM1, TNFRSF21, TTYH1, ZNF680, ZNF75A, JUND* |
| *SOX4* | *CFAP65; PINX1; SOX10; SOX11; SOX12; SOX13; SOX14; SOX15; SOX17; SOX18; SOX2; SOX3; SOX30; SOX4; SOX5; SOX6; SOX7; SOX8; SOX9; SRY* |  | *ABHD2, ASH1L, GFOD2, LAMP2, LYSMD3, MAGI3, OSBPL6, RIC3, SEL1L3, TNFRSF19, SOX4* |
| *ETS1* | *ILF2; NFATC2; PURA; GABPA; NKX2-1; GATA1; GATA2; GATA3; GATA4; GATA5; GATA6; ETS1; RARA; RARB; RARG; RXRA; RXRB; RXRG; THRA; THRB; YY2; STAT1; TCF3; SPI1; ELF3* | *ATF1; ATF2; ATF3; ATF4; ATF7; CREB1; CREM; CRX; E2F1; E2F6; EHF; ELF1; ELF2; ELF4; ELF5; ELK1; ELK3; ELK4; EP300; ERG; ETS1; ETS2; ETV2; ETV4; ETV6; ETV7; FLI1; FOS; FOSB; FOSL1; FOSL2; FOXG1; FOXO1; GABPA; GABPB1; GATA1; GATA2; GATA3; GATA4; GATA5; GATA6; HMGA1; HMGA2; HOXA13; IRF8; JUN; JUNB; JUND; KLF4; NFATC1; NFIA; NFIC; NKX2-1; NR3C1; SMAD4; SOX10; SOX4; SPI1; SPIB; STAT1; STAT6; TAF6; TBP; TCF7L1; TCF7L2; TFAP2C; YY1; ZEB1; ARID3A; CEBPB; ERF; HDX; ILF2; LEF1; NFATC2; NFATC3; NKX3-2; PAX4; PDX1; PURA; SIRT6; SRY; TAL1; TFAP4; ERF; RARA; RARB; RARG; RXRA; RXRB; RXRG; THRA; THRB; ETV1; GFI1; GFI1B; GLI3; HOXA13; IKZF1; ING4; LTF; MAFA; MZF1; OVOL2; POU3F1; ZBTB14; ZNF354C; SOX10; ZNF354C; ASCL1; CEBPA; CEBPD; CEBPE; CEBPG; CHURC1; HNF4A; HNF4G; IKZF2; MEIS1; MYB; MYCN; MYF5; MYF6; MYOD1; MYOG; NEUROD2; NR2F1; NR2F2; NR3C1; RFX1; RFX2; RFX3; RFX4; RFX5; RFXANK; RFXAP; SREBF1; SREBF2; TCF12; TCF4; TFAP4; YY2; FEV* | *ADAMTS7, ANGPTL2, CD34, CEBPB, PDGFB, SH2B3, SPRY4, ST6GALNAC6, VASP, ANGPT2, FAT1, FLT1, MGLL, PECAM1, PLXNA2, TIE1, TNFRSF1A, VAMP5, C1orf54, CAV1, CD93, EMCN, GJA1, GNG11, HES1, HLA-C, KCNE3, NOTCH4, PRSS23, RNF152, TFPI, AHNAK, APLN, CDH11, CDH5, CHST1, COL15A1, DLC1, DLL4, DYSF, EMILIN1, EMP1, EPAS1, ITGA5, JUP, MMRN2, MVP, PDE4B, PDE8A, PDLIM2, PGF, SERPINH1, SH2D3C, SHANK3, TRIB3, CAV2, ETS1, ITGA6, PNPO, SMOC1, ANTXR1, CLEC14A, ECSCR, EFEMP1, ESM1, HERPUD1, MYLIP, NFATC4, PLSCR1, TIMP3, TPM2, WFS1, OLFML2A, PLOD1, HSD11B2, PLS3* |
| *CEBPD* | *NFIA; NFIC; YY1* | *CEBPA; CEBPB; CEBPD; CEBPE; CEBPG; E2F1; FOXA1; FOXA2; FOXA3; HIC1; HLTF; HOXA13; IKZF2; MAFA; NANOG; NFE2L1; NR1H4; NR2F1; NR2F2; POU2F1; RXRA; SMAD3; SREBF2; STAT6; TCF7L1; TEAD1; TFAP2A; TFCP2; USF1; YY2; ZBTB14* | *CALD1, CTNND1, FHL2, HES1, HOPX, MBNL1, NFIB, POLD4, RASD2, TES, CEBPD* |
